# Genetic inactivation of the Translin/Trax RNase activity alters small RNAs including miRNAs, disrupts gene expression and impairs distinct forms of hippocampal synaptic plasticity and memory

**DOI:** 10.1101/2025.07.10.663777

**Authors:** Mahesh Shivarama Shetty, Junko Kasuya, Xiuping Fu, Marisol Carmela Lauffer, Satya Murthy Tadinada, Tania Chatterjee Chowdhury, Jay M. Baraban, Ted Abel

## Abstract

Neurons utilize RNA interference in the reversible translational repression of synaptically localized mRNAs, enabling rapid translation in response to synaptic activity. Two evolutionarily conserved proteins, Translin and Trax, form an RNase complex which processes miRNAs, tRNAs and siRNAs. To determine the specific role of the RNase activity of this complex in brain function, we employed a mouse line harboring a point mutation in Trax (E126A) that renders the Translin/Trax RNase inactive. At the molecular level, we found alterations in the levels of multiple small RNAs including miRNAs, tsRNAs and substantial downregulation of gene expression at the mRNA level in the hippocampus of TraxE126A mice. At the synaptic level, TraxE126A mice exhibit deficits in specific forms of long-term hippocampal synaptic plasticity. At the behavioral level, TraxE126A mice display impaired long-term spatial memory and altered open-field and acoustic-startle behavior. These studies reveal the functional role of Translin/Trax RNase in the mammalian brain.

## Introduction

Gene regulation plays a critical role in long-lasting forms of synaptic plasticity and long-term memory. Gene regulation occurs at multiple levels, including transcriptional, post-transcriptional, translational, post-translational and epigenetic regulation. Identifying the molecular players orchestrating these processes will help us better understand the mechanisms and develop strategies to target them. RNA interference (RNAi) is one such mechanism used by cells for post-transcriptional gene silencing. In RNAi, small interfering RNAs (siRNAs) produced by processing double-stranded RNA (dsRNA) are loaded into an effector complex called RNA-induced silencing complex (RISC) that contains target messenger RNA (mRNA), an RNase argonaute-2 (Ago2) and accessory proteins, which act to degrade or silence target mRNAs^1–3^. Similarly, microRNAs (miRNAs), upon cleavage from their precursor forms, also take part in post-transcriptional gene regulation by binding to the 3’-untranslated regions (3’-UTRs) of the target mRNAs thereby leading to their translational repression or degradation^4–8^. Neurons utilize this mechanism of miRNA-mediated translational repression to traffic and localize mRNAs in the vicinity of synapses where, in response to specific synaptic activity, miRNAs are rapidly degraded to allow translation of select set of mRNAs.^9–15^. Presence of numerous miRNAs and translationally silent mRNAs near the synapses supports this^9,16–18^. Several proteins involved in RNA trafficking, stabilization, compartmentalization, activity-mediated signaling and translational activation have been shown to regulate this process^15,18–21^.

The Translin/Trax complex (TN/TX), formed by Translin (also called TBRBP; testis, brain RNA-binding protein) and Trax (Translin-associated factor X) proteins, is an emerging player in the regulation of post-transcriptional gene expression^14,22–24^. Both Translin and Trax are evolutionarily conserved, possess nucleic acid binding capability and are implicated in several biological processes such as DNA damage repair^25,26^, homologous recombination^27,28^, cell division and proliferation^29,30^, genome stability^31^, spermatogenesis^32^, RNA trafficking and nucleic acid metabolism^33–35^. Apart from each other, they also have other interaction partners such as adenosine A2A receptor^36^, phospholipase C beta (PLCβ)^37^ and Glycogen synthase kinase 3 beta (GSK3β)^26^.

A key feature of the TN/TX complex is that it possesses endonuclease activity against single-stranded RNA (ssRNA), circular RNA and DNA^23,27,38,39^. The complex is also called C3PO (component 3 promoter of RISC) because in *Drosophila* and human systems it was found to enhance the activity of RISC by facilitating endoRNase cleavage of the siRNA passenger strand^23,24^. However, it was subsequently found that the RNase activity of the TN/TX complex can cleave many precursor and mature miRNAs by targeting mismatches in the stem region^14,15,39,40^. Studies from us and others have shown that the TN/TX complex indeed degrades select set of miRNAs to reverse the translational silencing of specific mRNA transcripts in hippocampal neurons^41^, in cerebellum^42^, in striatum^43,44^ and in vascular smooth muscle cells^45,46^. Thus, these findings suggest that the TN/TX complex appears to have opposite effects on silencing mediated by microRNAs and siRNAs. Furthermore, TN/TX RNase also plays a role in tRNA processing where it removes 5′ pre-tRNA fragments after the processing of pre-tRNAs by RNase P^34^. Growing evidence seems to suggest that small RNA fragments derived from tRNAs could have a role in translational regulation^47,48^.

Given the prominent expression of both Translin and Trax in the mammalian brain^40,49^, several studies have investigated their role in the nervous system and revealed that they are involved in mechanisms underlying synaptic plasticity and memory^14,41,50,51^, normal as well as addictive behavior^44,51,52^. Trax has also been proposed as a candidate susceptibility gene for psychiatric disorders^53–56^ and to provide neuroprotection in Huntington’s disease^43^. Important clues to the functional role of Translin and Trax proteins have been uncovered from studies using a mouse model that constitutively lacks Translin^41,50–52,57^. Interestingly, in the brain, constitutive deletion of Translin led to the degradation of Trax at the protein level^52^, highlighting a crucial role the TN/TX complex. Therefore, it is important to assess the specific role of TN/TX RNase activity in neuronal function without affecting other functions of Translin and Trax. Although the complex is heterooligomeric, comprising different ratios of Translin and Trax subunits^38^, the RNase activity of the complex is exclusively conferred by the Trax subunits and studies have identified three key acidic residues E126, E123 and D204 within Trax, each of which disrupts the catalytic activity when mutated^23^. Introduction of the E126A point mutation in Trax abolishes the RNase activity of the TN/TX complex against synthetic miRNA, miR-409, in an *in vitro* cleavage assay^42^. We have previously generated a mouse line that constitutively harbors the TraxE126A point mutation, which abolishes the RNase activity of the TN/TX complex without affecting protein expression, stability or the ability of Translin and Trax to interact with each other^42^. In this study, we utilized this point mutant mouse line to characterize the impact of impairing TN/TX RNase activity on various classes of small RNAs in the hippocampus and the subsequent alterations in synaptic plasticity, behavior, and long-term memory.

## Materials and Methods

All experiments were performed according to the National Institutes of Health guidelines and were fully approved by the Institutional Animal Care and Use Committee (IACUC) at the University of Iowa.

### TraxE126A point mutant mice

The generation and maintenance of TraxE126A point mutant mice were described previously^42^. Mice were on C57BL/6J background. Heterozygous male and female mice were mated to produce homozygous point mutant mice and wildtype littermates. Mice were maintained on a 12h light/12h dark cycle. Food and water were available *ad libitum*. Breeder mice were maintained on a diet with higher fat content (Teklad irradiated S-2335 mouse breeder diet #7904; 11.4% fat) to enhance breeding efficiency. Weaned mice were maintained on a standard mouse diet (Teklad NIH-31 irradiated modified mouse diet #7913; 6.2% fat). All experiments were performed during the light cycle using homozygous Trax point mutant mice and wildtype littermates as controls. Genotyping was performed either in-house using PCR or through service offered by Transnetyx (Cordova, TN).

### Small RNA sequencing

3-4 months old naïve male TraxE126A point mutant mice (n=4) and wildtype littermates (n=5) were sacrificed by cervical dislocation and hippocampi were quickly dissected in ice-cold, oxygenated aCSF. The hippocampi were collected in sterile, DNase and RNase-free tubes chilled on dry ice and immediately stored at –80 °C. Total RNA (including small RNAs) was isolated from these samples using Invitrogen miRVana miRNA Isolation Kit with phenol (Thermo Fisher Scientific, #AM1560) according to manufacturer’s instructions. The samples were treated with DNase I (Zymo, #E1010). The quality of the RNA samples was confirmed using NanoDrop 2000 Spectrophotometer and Agilent Bioanalyzer 2100. Small RNA cDNA libraries were prepared from these samples using Illumina TruSeq Small RNA Library Preparation Kit and were sequenced on one lane of the Illumina NovaSeq 6000 SP flowcell (50bp paired-end sequencing, ∼50 million pairs of reads) at the Iowa Institute of Human Genomics (IIHG) Core Facility at the University of Iowa.

<u>Data analysis</u>: Data analysis was performed using Partek Flow interface (Partek, Illumina, CA). Following QA/QC, 5’-TGGAATTCTCGGGTGCCAAGG-3’ adaptor sequence was trimmed from the 3’ end of read 1 and 5’-GATCGTCGGACTGTAGAACTCTGAAC-3’ adaptor sequence was removed from the 3’ end of read 2 using Cutadapt 4.2^58^, followed by base trimming. Trimmed reads were aligned using Bowtie^59^ and annotated using mm10 miRBase v22 mature miRNA database. The resulting miRNA counts were filtered to exclude miRNAs with minimum count <= 1 and normalized using median ratio method. The differential miRNA expression analysis was performed with DESeq2^60^. Differentially expressed miRNAs with P-values (adjusted for multiple comparison using Benjamini-Hochberg method) less than 0.050 and fold change of 1.150 were considered for functional enrichment analysis.

### Small RNA microarray

3-4 months old naïve male TraxE126A point mutant mice (n=4) and wildtype littermates (n=5) were sacrificed by cervical dislocation and hippocampi were quickly dissected in ice-cold, oxygenated aCSF. The hippocampi were collected in sterile, DNase and RNase-free tubes chilled on dry ice and immediately stored at –80 °C. Total RNA (including small RNAs) was isolated from these samples using Invitrogen miRVana miRNA Isolation Kit with phenol (Thermo Fisher Scientific, #AM1560) according to manufacturer’s instructions. The samples were treated with DNase I (Zymo, #E1010). The quality of the RNA samples was confirmed using NanoDrop 2000 Spectrophotometer and Agilent Bioanalyzer 2100. The samples were then shipped on dry ice to Arraystar Inc. (Rockville, Maryland) for simultaneously profiling major small RNA classes including miRNA, pre-miRNA, tsRNA, tRNA and snoRNA using Arraystar Mouse Small RNA Array V1.0. RNA sample quantity and quality control was performed again prior to microarray analysis using NanoDrop ND-1000 spectrophotometer and RNA integrity by Bioanalyzer 2100 or gel electrophoresis. The microarray workflow involves 3’ terminal dephosphorylation of total RNA (100 ng for each sample) with T4 polynucleotide kinase followed by DMSO denaturation and direct enzymatic 3’ end labeling with Cy3. The labeled RNA species are then hybridized onto Arraystar Small RNA Expression Microarray (8×15K format), scanned by an Agilent G2505C scanner followed by data processing and analysis. The Arraystar Mouse Small RNA Array V1.0 comprises a total of 14,192 distinct probes and includes the following coverage for different small RNA species: miRNAs – 1,949 (966 5-p and 983 3-p); pre-miRNAs – 1,122; tsRNAs – 1,767; mature tRNAs – 270; snoRNAs – 1,324. Following databases were used as sources for small RNA nomenclature and sequence: miRNA – miRBase(v22); tsRNA – tRFdb, GtRNAdb (Updated to 18.1 2019.08); pre-miRNA – miRBase(v22); tRNA – GtRNAdb(Updated to 18.1 2019.08), ENSEMBL(v99); snoRNA – ENSEMBL(v99).

#### Data Analysis

The intensity data were extracted from the acquired array images using Agilent Feature Extraction software (version 11.0.1.1). Quantile normalization and subsequent data processing were performed using GeneSpring GX v12.1 software package (Agilent Technologies). After normalization, the probe signals having Present (P) or Marginal (M) QC flags in at least 4 out of 9 samples were retained. Multiple probes from the same small RNA (miRNA/tsRNA(tRF&tiRNA)/pre-miRNA/tRNA/snoRNA) were combined into one RNA level. Differentially expressed small RNAs between two comparison groups were identified by fold change (FC; ≥1.150) and statistical significance (unpaired t-test P-value and False Discovery Rate <0.050) thresholds.

##### miRNA target prediction

The predicted mRNA targets of the selected miRNAs were acquired from the miRDB database^61^. Only the targets with Target Score ≥60 were included in the analysis.

### mRNA sequencing

3-4 months old naïve male TraxE126A point mutant mice (n=4) and wildtype littermates (n=5) were sacrificed by cervical dislocation and hippocampi were quickly dissected in ice-cold, oxygenated aCSF. The hippocampi were collected in sterile, DNase and RNase-free tubes chilled on dry ice and immediately stored at –80 °C. Total RNA (including small RNAs) was isolated from these samples using Invitrogen miRVana miRNA Isolation Kit with phenol (Thermo Fisher Scientific, #AM1560). The samples were treated with DNase I (Zymo, #E1010). The quality of the RNA samples was confirmed using NanoDrop 2000 Spectrophotometer and Agilent Bioanalyzer 2100. cDNA libraries were prepared from these samples using Illumina Stranded mRNA Library Preparation Kit with oligo-dT purification of polyadenylated RNA and were sequenced on one lane of the Illumina NovaSeq 6000 SP flowcell (50bp paired-end sequencing, ∼50 million pairs of reads) at the Iowa Institute of Human Genomics (IIHG) Core Facility at the University of Iowa.

#### Data analysis

Data analysis was performed using Partek Flow interface (Partek, Illumina, CA). Following QA/QC, raw reads were subjected to base trimming and aligned using STAR aligner version 2.7.8a^62^. Aligned reads were quantified by Partek E/M model and annotated to mouse reference genome mm39 Ensemble release 110. The resulting mRNA counts were filtered to exclude mRNAs with minimum count <= 1 and normalized using median ratio method. Differential gene expression analysis was performed with DESeq2^60^. Differentially expressed mRNAs with P-values (adjusted for multiple comparisons using Benjamini-Hochberg method) less than 0.050 and fold change of 1.200 were considered for functional enrichment analysis.

##### Functional annotation and enrichment

Functional annotation and pathway enrichment analysis was performed using DAVID platform^63,64^ with the EASE score threshold set at 0.05 and minimum gene count threshold set at 5. Kyoto Encyclopedia of Genes and Genomes (KEGG), Reactome and WikiPathways databases were used for functional enrichment of pathways. Gene Ontology (GO) and UniProt databases were used for enrichment analysis of GO terms. Pathways or GO terms with the Benjamini-Hochberg adjusted P-value threshold <0.050 were considered to be significantly enriched. Protein-Protein interaction clustering was performed using STRING database using k-means clustering method.

##### qRT-PCR

Total RNA was isolated from the hippocampi of WT and TraxE126A mutant mice (3-4 months old naïve males) using Invitrogen miRVana miRNA Isolation Kit with phenol (Thermo Fisher Scientific, #AM1560) according to manufacturer’s instructions. The samples were treated with DNase I (#E1010, Zymo Research, Irvine, CA) further purified using the Zymo RNA Clean & Concentrator-25 Kit (Cat. # R1017, Zymo Research). The quality of the RNA samples was confirmed using NanoDrop 2000 Spectrophotometer and Agilent Bioanalyzer 2100. Next, 1 µg of DNase-treated RNA was reverse-transcribed into cDNA using the iScript cDNA Synthesis Kit (Cat. # 1708890, Bio-Rad, Hercules, CA). Quantitative PCR (qPCR) was performed using SYBR™ Green Universal Master Mix (Cat. # 4309155, Thermo Fisher, Waltham, MA) in four technical replicates in three independent experiments across four and five biological replicates for each genotype. Primer sequences used for qRT-PCR are provided in Supplementary Table T1. The Δ threshold cycle (ΔC_T_) values were calculated by normalizing the C_T_ values of genes to those of Hprt. The relative fold change in the gene expression was determined using the formula: Fold Change = 2−ΔΔC_T_ against average value of WT samples.

### Electrophysiology

Experiments were performed as previously described^65,66^. Male and female mice (2.5-to 6-month-old for LTP experiments; 23-to 33-days-old for LTD experiments) were sacrificed by cervical dislocation and hippocampi were quickly collected in chilled, oxygenated artificial cerebrospinal fluid (aCSF; 124 mM NaCl, 4.4 mM KCl, 1.3 mM MgSO_4_⋅7H_2_O, 1 mM NaH_2_PO_4_⋅H_2_O, 26.2 mM NaHCO_3_, 2.5 mM CaCl_2_⋅2H_2_O and 10 mM D-glucose) bubbled with 95% O_2_ / 5% CO_2_. 400 μm-thick transverse slices were prepared from dorsal two-thirds of the hippocampus using a manual McIlwain slicer (Stoelting) and placed in an interface recording chamber at 29°C (Fine Science Tools, Foster City, CA). The slices were constantly perfused with aCSF at ∼1 ml/min. Slices were incubated for at least 2 hours in aCSF before starting the recordings. Field-EPSPs were recorded using aCSF-filled glass electrodes (resistance 2-5 MΩ) in the hippocampal area CA1 stratum radiatum by stimulating Schaffer collateral pathway with monopolar lacquer-coated stainless-steel electrodes (#571000; A-M Systems, Sequim, WA). Test stimulation was a biphasic, constant current pulse (0.1 ms duration per phase) delivered every minute at a stimulation intensity that evoked ∼40% (or ∼50% in LTD experiments) of the maximal fEPSP amplitude as determined by an input-output curve (stimulation intensity v/s fEPSP amplitude) in each experiment (stimulation intensity range 5 µA – 70 µA). Paired-pulse facilitation (PPF) was assessed by delivering a pair of stimuli at five different inter-stimulus intervals (ISI; 300 ms – 5 ms). A stable baseline response was recorded for at least 20 min before delivering LTP or LTD-inducing stimulus. LTP was induced by the following stimulation paradigms: spaced 4-train LTP paradigm comprised of four 100 Hz, 1 s trains (pulse width 0.1 ms per phase) delivered with an intertrain interval of 5 minutes; massed 4 train LTP paradigm comprised of four 100 Hz, 1 s trains (pulse width 0.1 ms per phase) delivered with an intertrain interval of 5 seconds; 1-train LTP stimulation comprised of a single 100 Hz train (1 s duration; pulse width 0.1 ms per phase). LTD was induced by 1 Hz stimulation for 15 minutes (900 pulses; pulse width 0.1 ms per phase). All stimulation protocols were delivered at the baseline stimulation intensity. In synaptic tagging and capture (STC) experiments, two stimulation electrodes were positioned in CA1 stratum radiatum on either side of a recording electrode to record fEPSPs evoked by the stimulation of two independent pathways (inputs). Pathway-independence was ensured by the lack of paired-pulse facilitation (at 50 ms ISI) between the two. Stimulus intensity was set to elicit ∼40% of the maximum field-EPSP amplitude determined by an input-output curve for each pathway. After a stable 20-min baseline response, massed 4-train stimulation was delivered to one pathway (S1) followed 30 min later by 1-train stimulation in the second pathway (S2).

#### Data Analysis

Data were acquired using Clampex 10 and Axon Digidata 1440/1550 digitizer (Molecular Devices, Union City, CA) at 20 kHz and were low-pass filtered at 2 kHz with a four-pole Bessel filter. Data analysis was performed using Clampfit 10 (Molecular Devices, Union City, CA). Data plots and statistical analyses were performed using GraphPad Prism10. Input-output and PPF data were analyzed using repeated measures ANOVA. The maintenance of LTP or LTD was analyzed by comparing the average of the normalized slopes over the last 20-min between two groups using unpaired t-test. The ‘n’ used in all the experiments represents the number of mice and replicate slices from the same mouse were averaged. Differences were considered statistically significant when P values were < 0.05. Data are plotted as mean ± S.E.M.

##### Behavioral Assays

Behavioral assays were performed using the facilities provided by the Neural Circuits and behavioral Core (NCBC) at the University of Iowa.

#### Open field

Male and female mice (2.5-to 5-month-old) were single-housed 4-5 days before the day of experiment and handled for 3-4 days (∼1-2 min every day) by the experimenter. On the day of the test, mice were introduced into the arenas (Dimensions: 38L x 32W x 30H) and allowed to explore the arenas freely for 6 minutes. The sessions were recorded by an overhead camera. The arenas were cleaned with 70% ethanol and allowed to dry prior to and between sessions. The videos were then analyzed using Ethovision XT program (Noldus) to score the time spent in the inner and outer areas (of equal area) of the arena, velocity and distance travelled. Data are reported as mean ± SEM. Statistical analysis was performed on GraphPad Prism 10 using unpaired t-tests.

#### Contextual fear conditioning (CFC)

Male and female mice (2.5-to 5-month-old) were single-housed 4-5 days before the day of experiment and handled for 3-4 days (∼1-2 min every day) by the experimenter. On the day of the training, each mouse was introduced to a context with a grid floor (FreezeScan System, CleverSys Inc. VA) and allowed to explore for 180 seconds during which a single electrical shock of 1.5 mA intensity, 2 s duration was delivered through the grid after 148 s. At the end of 180 s, mice were returned to their home cage and to the housing room. 24 hours after the training session, mice were reintroduced to the trained context for 5 minutes. Freezing behavior of the mice during the training and test sessions was scored using the FreezeScan software. ∼30-40 min after testing in the trained context, freezing was also assessed in an altered context with solid plastic floors and walls in the presence of a lemon soap odor. Average freezing levels before the shock (pre-shock) in the training session and during the 5 min test session in the trained and altered contexts were compared to measure context-specific freezing as a proxy for contextual fear memory. Data are reported as mean ± SEM. Statistical analysis was performed on GraphPad Prism 10 using two-way repeated measures ANOVA.

#### Acoustic Startle Response (ASR)

Acoustic startle responses were assessed using the SR-Lab System (San Diego Instruments) and the SR-Lab software, following previously described protocols^51,67,68^. Testing was performed in sound-attenuating chambers (35 x 33 x 38.5 cm), each equipped with a small ventilation fan and illuminated by a 15 W ceiling-mounted light bulb. Male and female mice (4-6-month-old) in their home cages were allowed to acclimate to the experimental room. Mice were restrained inside clear plexiglass cylinders (16 x 8.75 cm) positioned on plexiglass frames (12.5 x 20.5 x 0.6 cm), which were elevated 2.75 cm above a 30 x 30 x 4 cm base and habituated for 5 min with a 65dB background noise, all inside individual ASR chambers. Acoustic stimuli (65dB to 120dB, in multiples of 5dB; pseudorandom presentation) were delivered via a speaker located in the chamber ceiling, and startle responses were recorded using piezoelectric accelerometers mounted beneath each frame. Each startle stimulus was presented ten times in a pseudorandom order, with inter-trial intervals randomly set between 5 and 20 seconds. Signals were sampled at 1 kHz over a 65 ms period, with 10 ms immediately preceding stimulus onset designated as the baseline. These baseline values were used to screen for potential motion artifacts. Trials showing elevated or inconsistent baseline activity were excluded from analysis. Startle responses were averaged across the ten presentations at each stimulus intensity for each animal. Data are reported as mean ± SEM. Statistical analysis was performed on GraphPad Prism 10 using two-way repeated measures ANOVA.

#### Two-trial Y-maze

Group-housed male and female mice (2.5-to 5-month-old) were handled for 3-4 days (∼1-2 min every day) by the experimenter. On the day of the training, mouse was introduced to a Y-maze at the entry arm and allowed to explore the maze for 10 mins with one of the arms of the maze blocked. At the end of 10 min, mouse was returned to its home cage and the housing room. 24 hours after the training session, mouse was reintroduced to the Y-maze at the entry arm and allowed to explore the maze for 5 mins with none of the arms of the maze blocked. Distinct visual cues were present in each arm of the maze during both training and test sessions. The sessions were recorded using an overhead camera and the test session videos were manually scored using BORIS Software^69^ to assess the number of entries into the familiar and novel arms. Any time both the front paws of the mouse were past the location of the blocking slit in each arm, it was considered as an ‘entry’ into that arm. The time spent exploring the familiar and novel arms was also scored. For this, time spent exploring the novel and familiar arm including the respective halves of the central triangular area was included in the ‘exploration time’ for that arm (because the mice often tended to sit in the triangular area and sniff/explore the arm without fully entering the arm). Time spent grooming and time spent in the entry arm were not included. Data are presented as the discrimination index for the novel arm [(novel arm entry – familiar arm entry)/ total arm entries]. Data are reported as mean ± SEM. Statistical analysis was performed on GraphPad Prism 10 using unpaired t-tests.

### Figures and Illustration

Figures and illustration were generated using GraphPad Prism 10, Biorender and Adobe Illustrator. Upset Plot for the miRNA targets was generated using Express Analyst^70^. Venn Diagrams were generated using an online tool hosted by the ‘Good Calculators’ webpage.

### Data availability

The miRNA and mRNA sequencing datasets generated in this study have been deposited at the Gene Expression Omnibus (GEO) database with the accession number GSE300964.

## Results

### Expression of multiple small RNA species including miRNAs is altered in the hippocampus of mice carrying RNase-dead Trax

We first assessed the impact of abolishing the RNase activity of the TN/TX complex on the expression of small RNAs in the hippocampus of TraxE126A point mutant mice^42^. Based on our previous work with TN/TX ^41^, our primary interest was in examining role of TN/TX RNase on the expression of miRNAs in the hippocampus. We performed miRNA sequencing using total RNA extracted from the hippocampus of naïve adult male mice and found that the expression of multiple mature miRNAs was significantly altered in the TraxE126A mutants compared to their wildtype littermates (Fig. 1A and **Supplementary File 1**). Previous studies have shown that the TN/TX complex is involved in the degradation of precursor miRNAs (pre-miRNAs)^15,39,46,71^. However, we were not able to assess the alterations in pre-miRNA levels from our small RNA sequencing data because the kit used for cDNA library preparation was based on Dicer recognition sequences. Therefore, we analyzed the same total RNA samples using a small RNA microarray, which utilizes direct end labeling and allows us to determine the expression of multiple small RNA species including mature miRNAs, pre-miRNAs, small nucleolar RNAs (snoRNAs), mature transfer RNAs (tRNAs) and tRNA-derived small RNAs (tsRNAs). Microarray analysis also revealed that the expression of multiple mature miRNAs was altered in the hippocampus of TraxE126A mutants (Fig. 1B and **Supplementary File 2**). Twelve miRNAs (10 upregulated: miR-412-5p, miR-496a-3p, miR-409-3p, miR-409-5p, miR-330-3p, miR-330-5p, miR-128-2-5p, miR-667-3p, miR-378a-5p, miR-125b-2-3p; 2 downregulated: miR-3068-3p and miR-143-3p) were identified both in the miRNA sequencing and in the microarray analysis. Surprisingly, microarray analysis identified no pre-miRNA to be significantly upregulated and only one pre-miRNA (pre-miR-378b) to be significantly downregulated in the hippocampus of TraxE126A mutants (Fig. 1C and **Supplementary File 3**). Among snoRNAs, only two (Mir1949-201 and Gm24598-201) were found to be significantly upregulated in the TraxE126A mutants (Fig. 1D and **Supplementary File 3**). Microarray analysis of mature tRNAs identified five tRNAs to be significantly upregulated in the TraxE126A mutants and none were found to be downregulated (Fig. 1E and **Supplementary File 3**). The largest alteration among the small RNA species investigated using microarray analysis was found in the tsRNA species with 91 tsRNAs significantly upregulated and 22 tsRNAs significantly downregulated in the hippocampus of TraxE126A mutants (Fig. 1F and **Supplementary File 3**). Interestingly, almost all the significantly altered tsRNAs were 5’-fragments or halves.

**Figure 1.**
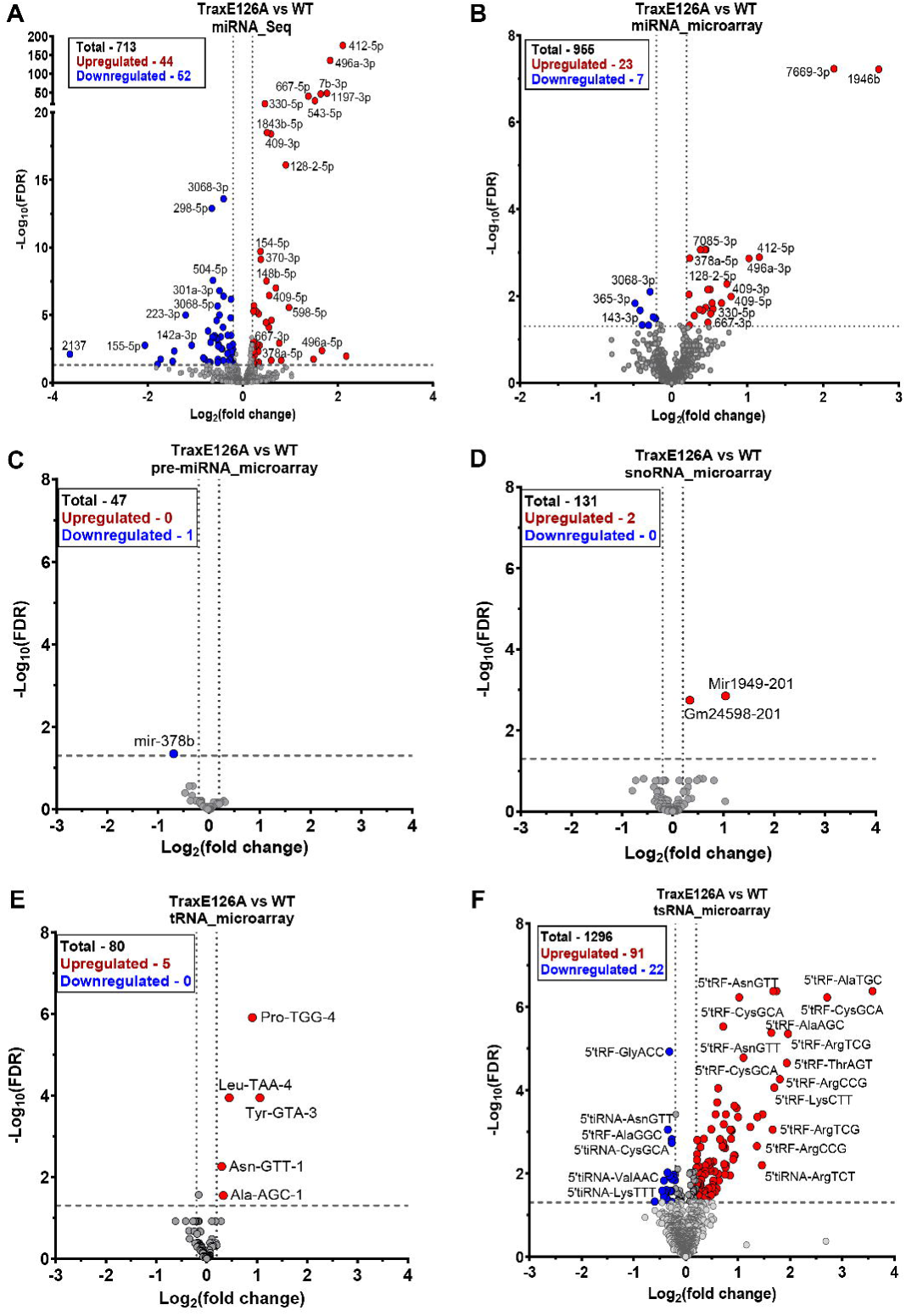
Expression of many small RNA species including mature microRNAs is altered in the hippocampus of TraxE126A mutant mice. Volcano plots showing the mature miRNAs differentially expressed in the TraxE126A mutants compared to wildtype littermates as identified by (A) miRNA sequencing and (B) miRNA microarray. Volcano plots showing the differentially expressed small RNA species identified using microarray analysis including (C) Precursor miRNAs (pre-miRNAs), (D) Small nucleolar RNAs (snoRNAs), (E) mature tRNAs and (F) tRNA-derived small RNAs (tsRNAs). In all the plots, the upregulated and downregulated miRNAs (false discovery rate, FDR <0.050 and log_2_fold change ≥0.200) are highlighted in red and blue, respectively. (TraxE126A, n=4; WT, n=5, all males). Largest changes were seen in tsRNA levels (majority are 5’-fragments) and mature miRNAs.

These results suggest that the RNase activity of the TN/TX complex could be involved in the regulation of multiple small RNA species, especially miRNAs and tsRNAs.

### MicroRNAs regulated by TN/TX RNase activity target transcripts predicted to be involved in synaptic plasticity, neuronal function and cell adhesion

Previous studies have provided evidence to show that TN/TX complex is involved in the degradation of miRNAs and thereby relieves the translational repression of the mRNA targets of those miRNAs in multiple tissues including brain^15,39,42,46,71^. Our own work has shown that global deletion of Translin and Trax leads to impairments hippocampus-dependent memory and synaptic plasticity^41,50^, at least in part due to accumulation of a set of miRNAs which repress the activity-induced translation of activin receptor type 1C (Acvr1c) mRNA^41^. Therefore, we focused our further investigation in this study on miRNAs.

We compared the mature miRNAs identified to be significantly altered in our small RNA sequencing analysis (Fig. 1A) and the microarray analysis (Fig. 1B). This revealed 12 common miRNAs, including 10 upregulated (miR-412-5p, miR-496a-3p, miR-409-3p, miR-409-5p, miR-330-3p, miR-330-5p, miR-128-2-5p, miR-667-3p, miR-378a-5p and miR-125b-2-3p) and 2 downregulated ones (miR-3068-3p and miR-143-3p), that were identified in both the analyses (Fig. 2A). Because of the greater confidence in these miRNAs as being regulated by TN/TX RNase activity, in this study, we focused on these 12 common miRNAs for downstream target prediction. We used the public miRNA target prediction database miRDB^61^ to identify mRNAs predicted to be targeted by the 12 common miRNAs. We only included targets with miRDB Target Score ≥60, indicating higher confidence. Although there were some shared targets between different miRNAs, a significant proportion of the targets were distinct for each miRNA (Fig. 2B and **Supplementary File 4**). We then used the combined list of predicted targets (miRDB Target Score ≥ 60; total 2921 targets) of all the 12 common miRNAs to perform functional enrichment and annotation analysis using the DAVID database^63,64^. Top KEGG pathways that showed significant enrichment (ranked by adjusted P-value) included MAPK signaling, axon guidance, long-term depression, PI3K-Akt signaling, focal adhesion, Ras signaling pathway among others (Fig. 2C and **Supplementary File 4**). Top Gene Ontology (GO) Biological Process terms that showed significant enrichment (ranked by adjusted P-value) included regulation of transcription by RNA polymerase II, nervous system development and cell adhesion among others (Fig. 2D and **Supplementary File 4**). We validated the expression levels of some of the predicted targets with prior evidence for involvement in neuronal function and plasticity using qRT-PCR and found that only some of these selected targets were differentially expressed in the hippocampus of TraxE126A mice compared to WT (**Supplementary Fig. S1**).

**Figure 2.**
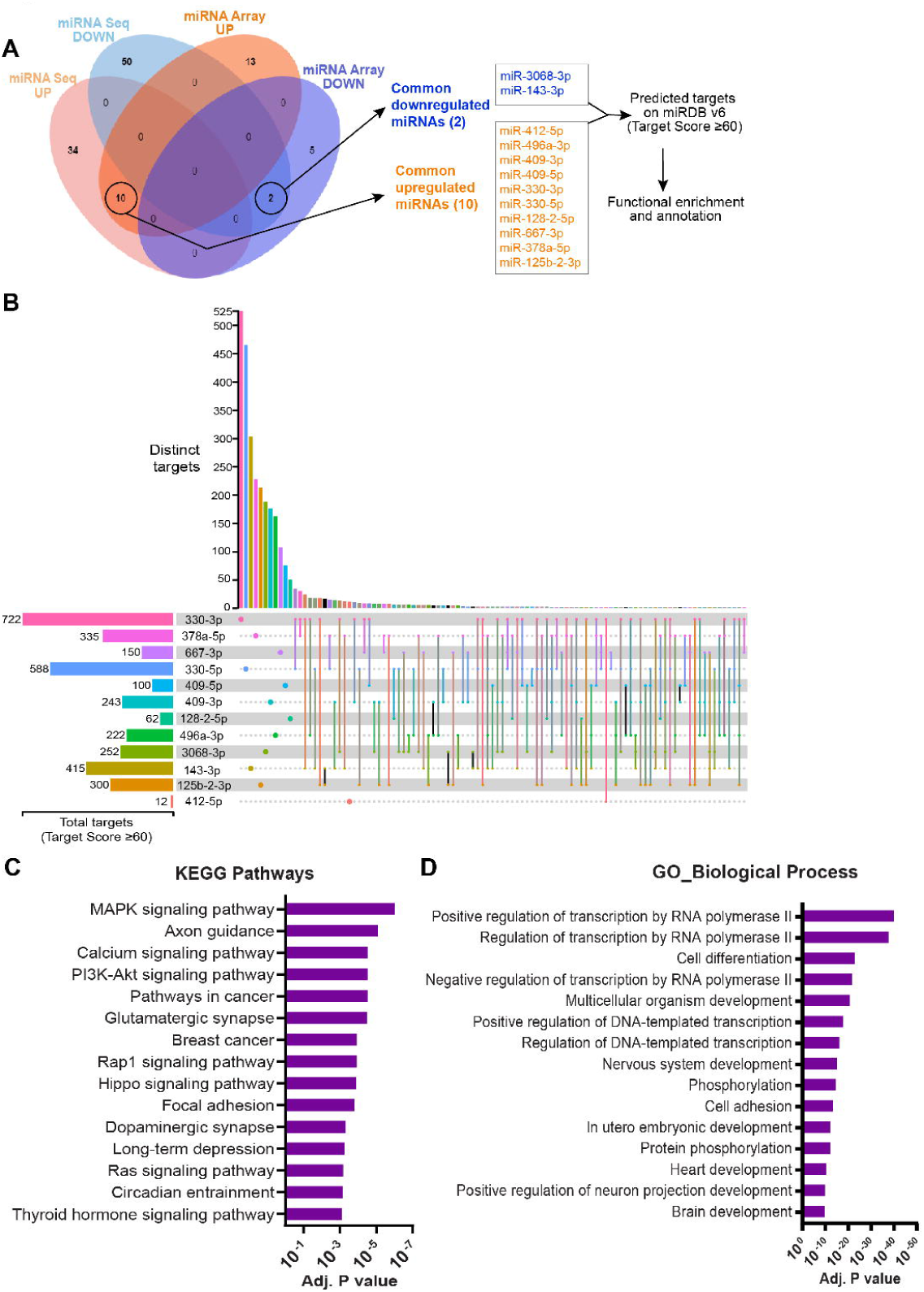
The predicted targets of a subset of mature miRNAs differentially expressed in the hippocampus of TraxE126A mutants show functional enrichment for pathways involved in neuronal function and plasticity. (A) Venn diagram showing the overlap between mature miRNAs identified using miRNA sequencing and microarray analysis (with FDR<0.050 and log_2_fold change ≥0.200). A total of 12 miRNAs (10 upregulated and 2 downregulated) were found to be common and were used for target prediction using miRDB database. (B) An upset plot showing the shared and unique predicted mRNA target profiles in the miRDB database for the 12 common miRNAs. Only targets with miRDB Target Score ≥60 are included. (C) Top 15 KEGG pathways and (D) Gene Ontology (GO) Biological Process terms from the functional enrichment analysis of the predicted targets of the 12 common miRNAs performed using DAVID database.

### mRNAs regulated by TN/TX RNase activity show enrichment for pathways involved in immune response, extracellular matrix signaling and cell adhesion

Because it is difficult to individually validate the huge number of predicted targets of the selected miRNAs, next, we performed mRNA sequencing from the same total RNA samples that were used for small RNA sequencing and microarray analysis. We reasoned that this would help us identify the overlap between the predicted targets of the miRNAs and changes in mRNA expression levels. The mRNA sequencing analysis revealed alterations in the expression of a significant number of genes with a large proportion of them being downregulated (Fig. 3A and **Supplementary File 5**). We then performed functional pathway enrichment analysis on the differentially expressed genes using DAVID database. Top KEGG pathways that showed significant enrichment (ranked by adjusted P-value) included ECM-receptor interaction, response to viral infection, focal adhesion, PI3K-Akt signaling pathway, complement and coagulation cascades, cell adhesion among others (Fig. 3B and **Supplementary File 5**). Top GO-Biological Process terms that showed significant enrichment (ranked by adjusted P-value) included immune response and related signaling, cell adhesion, integrin-mediated signaling, extracellular matrix organization among others (Fig. 3C and **Supplementary File 5**). Top GO-Molecular Function terms that showed significant enrichment (ranked by adjusted P-value) included protein binding, integrin binding, extracellular matrix components, nucleotide and ATP-binding and kinase activity among others (Fig. 3D and **Supplementary File 5**).

**Figure 3.**
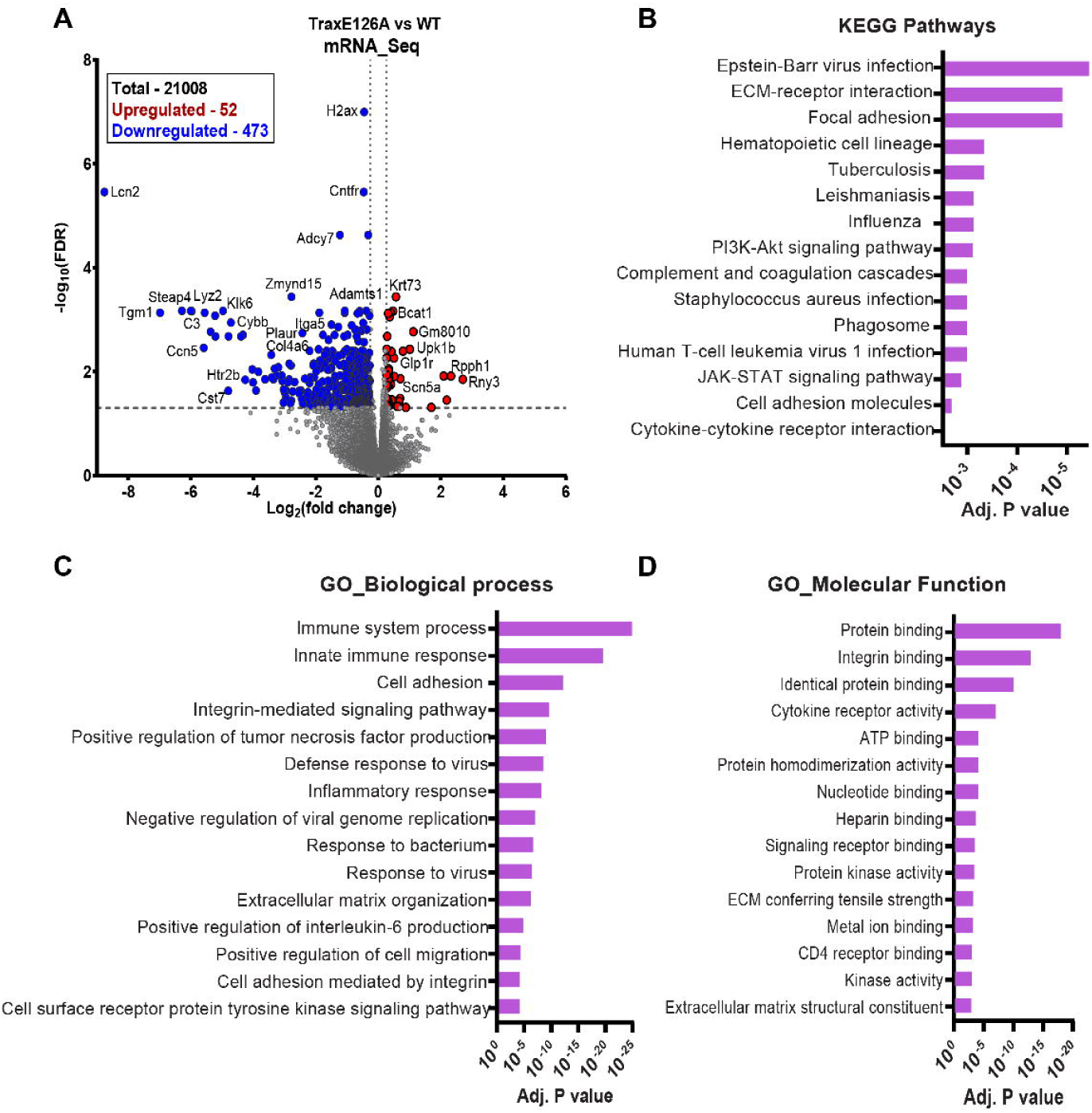
Abolishing the RNase activity of Trax leads to significant alterations in the expression of genes involved in immune function, extracellular matrix, and integrin-mediated cell signaling in the hippocampus. (A) Volcano plot showing the differentially expressed mRNAs in the hippocampus of TraxE126A mutants (n=4) compared to wildtype littermates (n=5). The upregulated and downregulated mRNAs (FDR<0.050 and log_2_fold change ≥0.263) are highlighted in red and blue, respectively. (B) Top 15 KEGG pathways from the functional enrichment analysis of the differentially expressed mRNAs performed using DAVID database. (C) Top 15 Gene Ontology (GO) Biological Process and (D) Molecular Function terms from the functional enrichment analysis of the differentially expressed mRNAs performed using DAVID database.

Together, these results indicate that abolishing the RNase activity of the TN/TX complex leads to significant alterations in the expression levels of genes involved in many signaling pathways such as extracellular matrix-mediated signaling, focal adhesion, cell adhesion, PI3K-Akt signaling and immune response. Some of these pathways were also annotated to be regulated by the predicted gene targets of 12 miRNAs altered in the TraxE126A mice (see Fig. 2C and **Supplementary File 4**).

### miRNAs targeted by TN/TX RNase regulate the expression of genes involved in focal adhesion, integrin signaling, and extracellular matrix in the hippocampus

To identify the extent of overlap between the predicted targets (2921) of the 12 common differentially expressed miRNAs and the mRNAs (525) that we identified as significantly altered in the TraxE126A mutants, we compared these lists (Fig. 4A). Surprisingly, only a small number of the predicted targets (63 out of 2921) were identified as significantly altered in the mRNA sequencing analysis. Our hypothesis was that miRNAs generally act to inhibit the expression of genes and therefore, we predicted an inverse relationship between the miRNA levels and their target mRNA levels. Accordingly, we were specifically interested in the targets of upregulated miRNAs which are found to be downregulated in mRNA sequencing analysis, and the targets of downregulated miRNAs which are found to be upregulated in mRNA sequencing analysis. We found 43 of the predicted targets of the upregulated miRNAs to be significantly downregulated at mRNA levels whereas none of the predicted targets of the downregulated miRNAs to be significantly upregulated at mRNA levels (Fig. 4A and **Supplementary File 5**). We focused on these 43 genes (Fig. 4A **inset**) and performed functional enrichment analysis using DAVID. Focal adhesion was the only pathway that showed significant enrichment in KEGG and WikiPathways databases (Fig. 4B). GO term analysis identified integrin binding and protein binding as the enriched Molecular Function terms, and focal adhesion, cell surface and collagen-containing extracellular matrix as the enriched Cellular Component terms. The GO-Biological Process terms that showed highest enrichment included cell adhesion, integrin-mediated signaling, immune response pathways and receptor tyrosine kinase signaling among others (Fig. 4B). We also performed clustering analysis on these 43 targets based on protein-protein interaction data using STRING database which identified significantly enriched clustering of genes involved in focal adhesion, extracellular matrix and integrin signaling (Fig. 4C). Next, we further validated the expression of a selected few genes involved in these signaling pathways at the mRNA level using qRT-PCR and confirmed that they are indeed downregulated (Fig. 4D).

**Figure 4.**
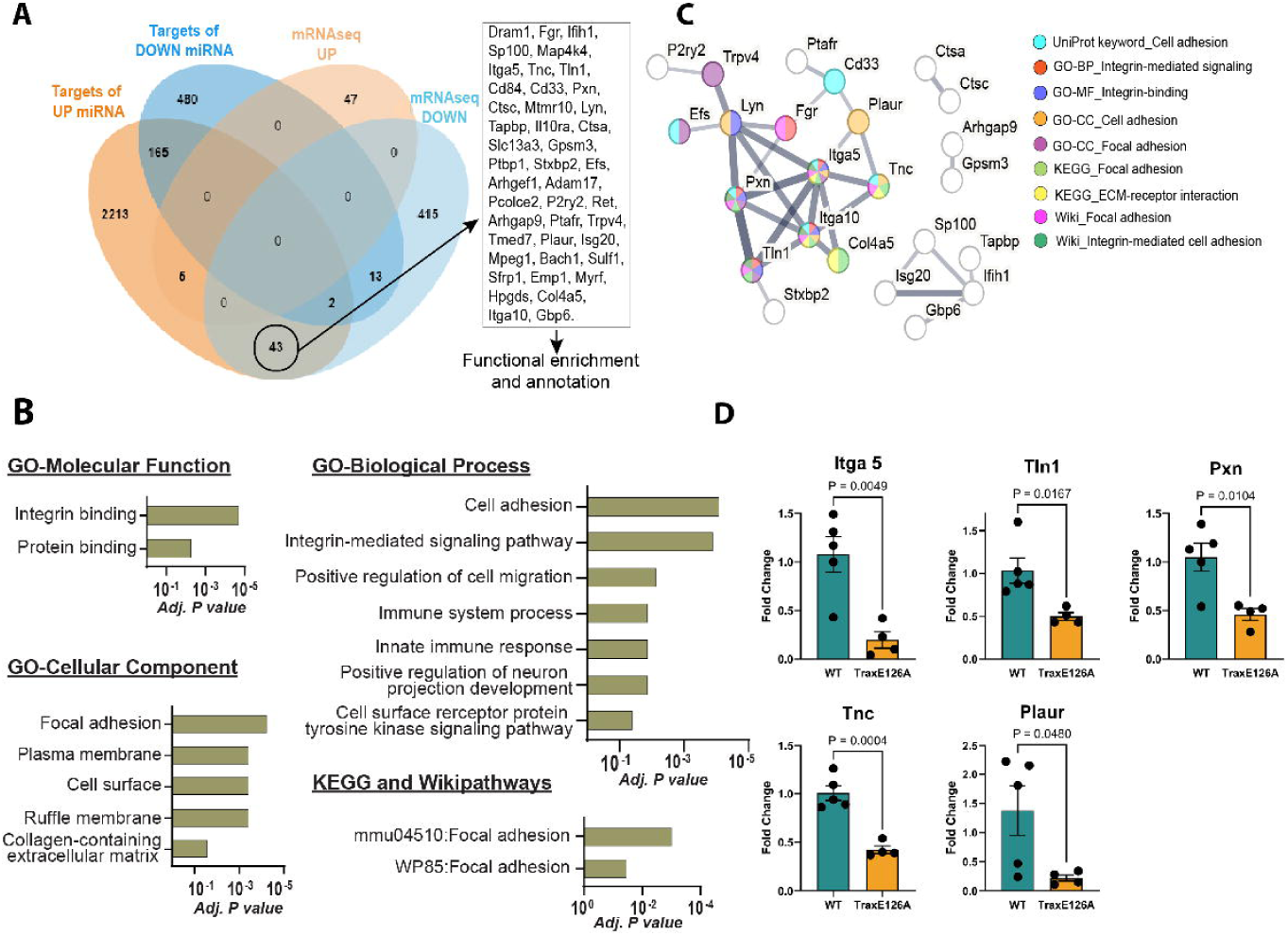
Components of focal adhesion, integrin signaling, and extracellular matrix are significantly downregulated in the hippocampus of TraxE126A mutant mice. **(A)** Venn diagram showing the overlap between the predicted targets (miRDB Target Score ≥60) of the 12 common miRNAs upregulated (UP miRNAs) or downregulated (DOWN miRNAs) (FDR<0.050 and log_2_fold change ≥0.200) and mRNAs upregulated (mRNAseq UP) or downregulated (mRNAseq DOWN) (FDR<0.050 and log_2_fold change ≥0.263) in the TraxE126A mutants. Inset shows the 43 genes that are targets of upregulated miRNAs and are also found to be downregulated in mRNA-seq. No genes were found that were targets of downregulated miRNAs and were upregulated in the mutants. **(B)** Significantly enriched (adjusted P-value <0.050) pathways (KEGG and WikiPathways) and Gene Ontology (GO) terms from the functional enrichment analysis of the 43 downregulated genes that are also targets of the upregulated miRNAs performed using DAVID database. **(C)** Clustering analysis of these 43 genes using STRING protein-protein interaction database showing clustering of genes involved in focal adhesion, integrin signaling and extracellular matrix. Genes within the cluster are also colored for significant functional enrichment of these terms in GO, KEGG and WikiPathways databases. **(D)** qPCR validation of the expression of selected genes (Integrin alpha 5, Talin1, Paxilin, Tenascin C and Plasminogen activator urokinase receptor) involved in focal adhesion, integrin signaling and extracellular matrix showed significant downregulation in the hippocampus of TraxE126A mice (n=4, males) compared to WT mice (n=5, males).

Together, these results indicate that miRNAs targeted for degradation by TN/TX RNase regulate the expression of genes involved in focal adhesion, integrin signaling, and extracellular matrix in the hippocampus.

### Abolishing the RNase activity of the TN/TX complex leads to impairments in specific forms of hippocampal synaptic plasticity and synaptic tagging/capture

Previously, we have shown that global deletion of Translin and Trax proteins leads to impairments in distinct forms of synaptic plasticity in the hippocampal CA1 region^41,50^. However, both Translin and Trax have been shown to have independent functions in addition to their role as a proposed miRNA-degrading complex^14,23,25,26,28,33,34,37,40,49^. Because the TraxE126A point mutation only abolishes the RNase activity of the complex but affects neither the protein levels of Translin and Trax nor their interaction in the hippocampus^42^, we assessed its impact on hippocampal synaptic plasticity to determine the specific role of the RNase activity of the TN/TX complex. Using acute hippocampal slices from WT and TraxE126A mutant mice (both males and females), we performed field-EPSP recordings in the CA1 stratum radiatum region by stimulating the Schaffer collateral pathway from CA3. We first assessed basal synaptic transmission using input-output responses and found no changes in the presynaptic fiber volley or fEPSP amplitudes (**Supplementary Fig.S2A and S2B**). We also measured paired-pulse facilitation (PPF), a very short-term plasticity, and found no significant alterations over a range of inter-stimulus intervals (**Supplementary Fig.S2C**). We then tested synaptic plasticity in response to distinct stimulation paradigms that induce short-term or long-term LTP and LTD. We found that the slices from TraxE126A mutant mice showed deficits only in the long-lasting LTP induced by spaced 4-train stimulation paradigm (Fig. 5A**-B**) but not by massed 4-train stimulation paradigm (Fig. 5C**-D**). TraxE126A mutant mice showed no impairments either in a short-lasting form of LTP induced by 1-train stimulation (Fig. 5E**-F**) or in LTD induced by prolonged low-frequency stimulation (Fig. 5G**-H**).

**Figure 5.**
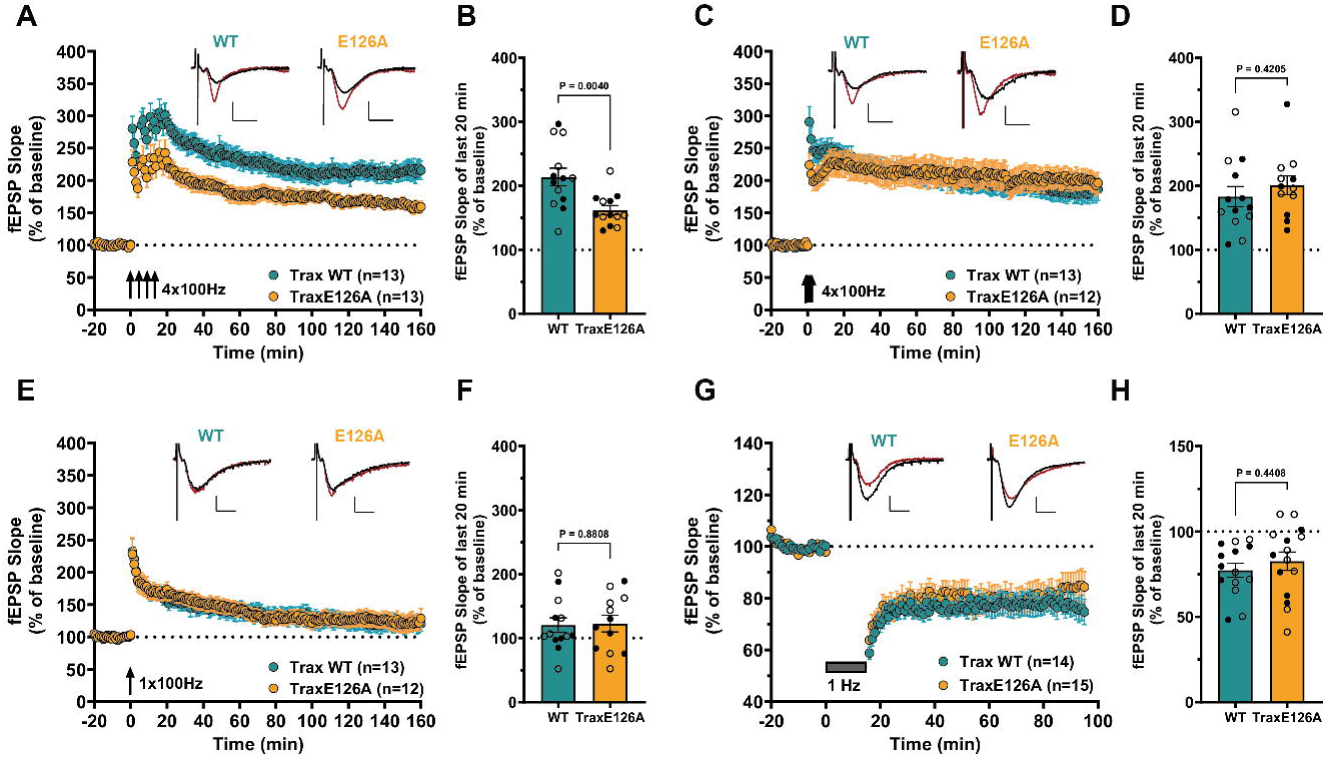
TraxE126A mutant mice show deficits in distinct forms of hippocampal synaptic plasticity in the hippocampal CA1 region. (A)-(B) Long-lasting LTP induced by spaced-4-train stimulation protocol is impaired in slices from TraxE126A mutant mice (n=13 mice, 20 slices; 7 males, 6 females) compared to WT littermates (n=13 mice, 17 slices; 6 males, 7 females). The mean fEPSP slope over the last 20 min recordings is significantly lower in TraxE126A mutants compared to WT (Welch’s t-test, two-tailed, t=3.307, df=17.76, P=0.0040, eta squared=0.381). **(C)-(D)** Long-lasting LTP induced by massed-4-train stimulation protocol is not impaired in slices from TraxE126A mutant mice (n=12 mice, 15 slices; 6 males, 6 females) compared to WT littermates (n=13 mice, 16 slices; 7 males, 6 females). The mean fEPSP slope over the last 20 min recordings is not significantly different in TraxE126A mutants compared to WT (Unpaired t-test, two-tailed, t=0.820, df=23, P=0.4205, eta squared = 0.028). **(E)-(F)** Short-lasting LTP induced by 1-train stimulation protocol is identical in slices from TraxE126A mutant mice (n=12 mice, 14 slices; 6 males, 6 females) compared to WT littermates (n=13 mice, 14 slices; 6 males, 7 females). The mean fEPSP slope over the last 20 min recordings is not significantly different in TraxE126A mutants compared to WT (Unpaired t-test, two-tailed, t=0.152, df=23, P=0.8808, eta squared = 0.0009). **(G)-(H)** LTD induced by 1 Hz, 15 min stimulation protocol shows no significant alterations in slices from TraxE126A mutant mice (n=15 mice, 24 slices; 6 males, 9 females) compared to WT littermates (n=14 mice, 24 slices; 7 males, 7 females). The mean fEPSP slope over the last 20 min recordings is not significantly different in TraxE126A mutants compared to WT (Unpaired t-test, two-tailed, t=0.782, df=27, P=0.4408, eta squared = 0.022). In all bar graphs, open symbols represent females and closed symbols represent males. Representative fEPSP traces are shown in the insets for each genotype, one sampled at baseline (black trace) and another at the end of the recording (red trace). Calibration bars for fEPSP traces: 2 mV vertical, 5 ms horizontal. The error bars represent SEM in all graphs.

We next investigated synaptic tagging and capture (STC), a form of associative heterosynaptic plasticity^72–74^. STC involves the interaction between local synaptic events (the ‘tag’) and broader dendritic or somatic events, such as protein synthesis, that supply the plasticity-related products (PRPs) for synapse-strengthening (‘capture’)^73,74^. This enables the heterosynaptic association between two synaptic plasticity events (e.g., LTP or LTD) that occur within a specific temporal window in synaptic inputs onto a common population of neurons. We have previously found that STC is impaired in the hippocampal slices from mice globally lacking both Translin and Trax proteins^41^. To know whether this impairment is specifically due to the lack of the RNase activity of TN/TX complex, we investigated STC in the hippocampal slices from TraxE126A mutant and WT mice using a similar two-pathway design to stimulate two independent synaptic inputs and recorded fEPSPs in the CA1 stratum radiatum (Fig. 6A). In each experiment, we confirmed that the two pathways were independent using lack of PPF (50 ms ISI) between the pathways as a measure. After 20 min of baseline recording, massed 4-train stimulation was delivered to one pathway (S1) followed 30 min later by 1-train stimulation in the second pathway (S2) (Fig. 6B). In the slices from WT mice, 1-train stimulation in S2 results in long-lasting LTP thereby demonstrating intact STC (Fig. 6C) whereas in slices from TraxE126A mutant mice, this heterosynaptic facilitation of 1-train LTP into a long-lasting LTP is impaired (Fig. 6D). Upon comparison of LTP maintenance between genotypes, massed 4-train LTP in pathway S1 showed no significant difference (Fig. 6E) whereas 1-train LTP in pathway S2 was significantly lower in mutants than WT (Fig. 6F).

**Figure 6.**
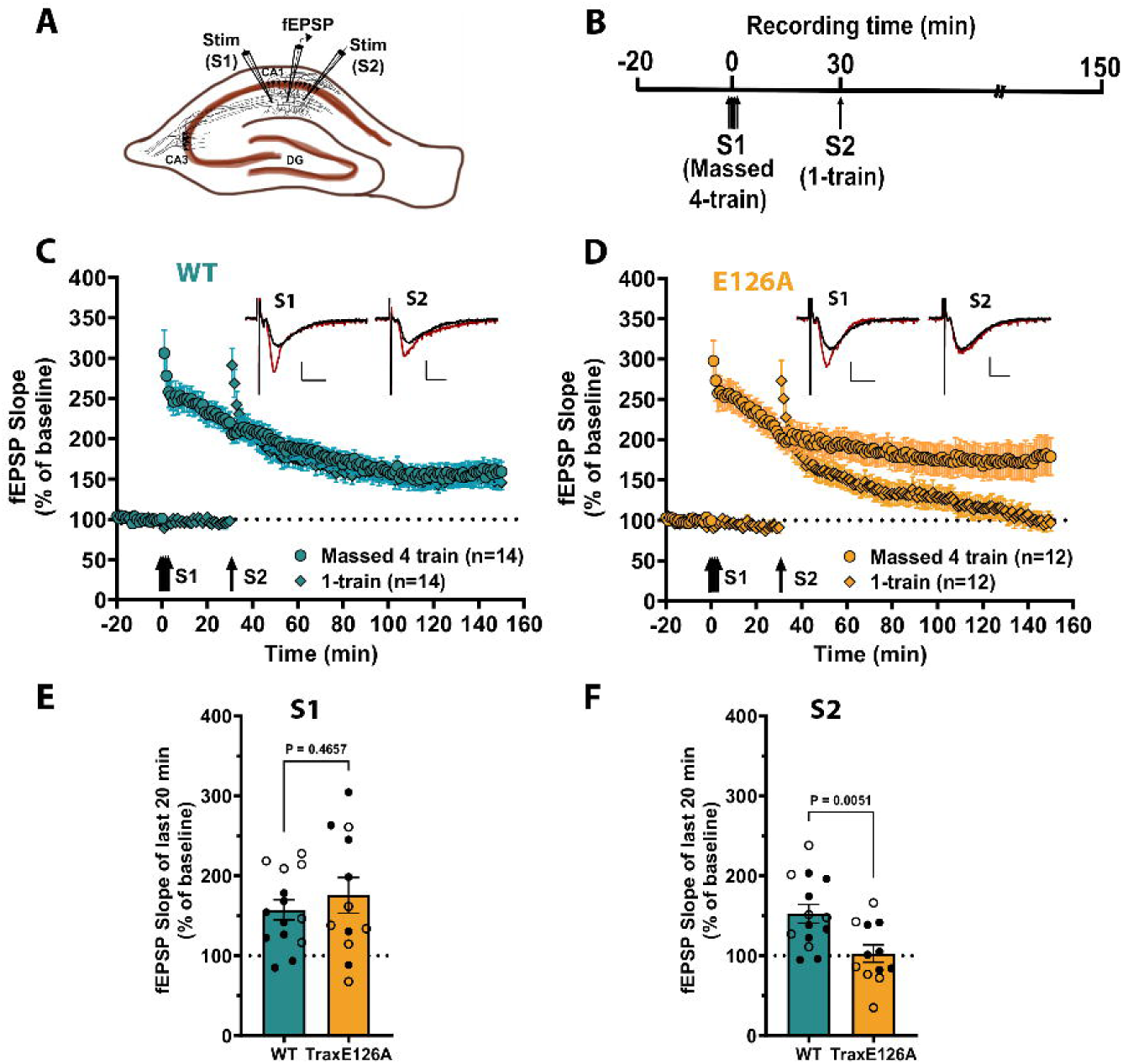
TraxE126A mutant mice show deficits in synaptic tagging and capture, a form of associative synaptic plasticity, in the hippocampal CA1 region. **(A)** A schematic representation of a hippocampal slice showing the location of electrodes in the hippocampal CA1 stratum radiatum. Two electrodes S1 and S2 are used to stimulate two independent Schaffer collateral inputs onto a population of neurons and the fEPSP responses are recorded with a common recording electrode. **(B)** Timeline of the synaptic tagging and capture (STC) experiments. After a stable baseline recording of 20 min in both the inputs, massed 4-train stimulation is delivered to input S1 followed 30 min later by 1-train stimulation in input S2. fEPSPs in both inputs are recorded for 2 hours after the 1-train stimulation. **(C)** STC in the slices of WT mice (n=14 mice, 16 slices; 8 males, 6 females) as shown by the successful facilitation of short-lasting LTP to long-lasting LTP in input S2 in response to weak 1-train stimulation. **(D)** In the slices of TraxE126A mutant mice (n=12 mice, 13 slices; 6 males, 6 females), 1-train stimulation in S2 only results in short-lasting LTP, demonstrating impaired STC. **(E)** A bar graph of the mean fEPSP slope over the last 20 min recordings in input S1 (massed 4-train) shows no significant difference in potentiation between TraxE126A mutants and WT (Unpaired t-test, two-tailed, t=0.741, df=24, P=0.4657, eta squared = 0.022). **(F)** A bar graph of the mean fEPSP slope over the last 20 min recordings in input S2 (1-train) shows a significantly lower potentiation in TraxE126A mutant slices compared to WT; (Unpaired t-test, two-tailed, t=3.080, df=24, P=0.0051, eta squared = 0.283). In all bar graphs, open symbols represent females and closed symbols represent males. Representative fEPSP traces are shown in the insets for each input at baseline (black trace) and at the end of the recording (red trace). Calibration bars for traces: 2 mV vertical, 5 ms horizontal. The error bars represent SEM in all graphs.

Together, these results show that TraxE126A mice which lack the RNase activity of the TN/TX complex exhibit deficits only in specific forms of hippocampal synaptic plasticity such as spaced 4-train LTP and STC, which phenocopy the synaptic plasticity deficits in global Translin KO mice^41,50^.

### Mice lacking the RNase activity of TN/TX complex show alterations in open-field behavior, acoustic startle response and impaired long-term memory in a spatial Y-maze task

We next tested the impact of abolishing the RNase activity of TN/TX complex on certain behavioral measures and long-term memory. We first checked open-field behavior because alterations in this could confound the use of spatial object exploration tasks for long-term memory assessment. To assess open-field behavior, WT and TraxE126A mice (both males and females) were allowed to individually explore an open arena for 6 min and time spent in the central and peripheral areas, total distance travelled, and mean velocity were calculated. TraxE126A mutant mice spent significantly lesser time in the central area of the arena, travelled significantly lesser distance and moved at a significantly lesser mean velocity compared to WT mice (Fig. 7A). We then assessed long-term contextual fear memory in these mice where they were allowed to explore a grid-floor context for 3 minutes during which they received a single foot shock (1.5mA, 2s). After 24 hours, mice were re-introduced to the same context for 5 minutes and their freezing levels were measured. Subsequently, freezing levels were also tested in an altered context. Increased freezing in the shocked context compared to pre-shock levels was considered as a proxy for long-term fear memory. The data revealed no significant alteration in contextual fear memory in TraxE126A mice compared to WT mice (Fig.7B). We next measured acoustic startle responses in these mice, which helps assess anxiety levels and sensorimotor processing. Mice were enclosed in an acoustic chamber and their startle responses were measured in response to the presentation of a series of acoustic stimuli ranging 65-120 dB. We found that when compared to the WT mice, TraxE126A mutant mice showed significantly dampened startle response to acoustic stimuli over 100 dB intensity (Fig.7C). Finally, we performed a spatial version of the Y-maze task involving a single training session and a test session (Fig.7D). During training session (10 min), mice were allowed to explore the Y-maze with one of its arms blocked. After 24 hours, mice were re-introduced to the maze and allowed to explore all the three arms (5 min test session) during which the number of entries into the novel and familiar arms as well as the time spent exploring each of the arms were measured. The WT mice made significantly more entries into the novel arm during the test session whereas TraxE126A mice did not show this preference for the novel arm demonstrating a deficit in long-term memory. Intriguingly, the time spent exploring the novel arm was not different between WT and TraxE126A mice (**Supplementary Fig. S3**), which could partly be due to the criteria used for scoring exploration time (**Supplementary Fig. S4**).

**Figure 7.**
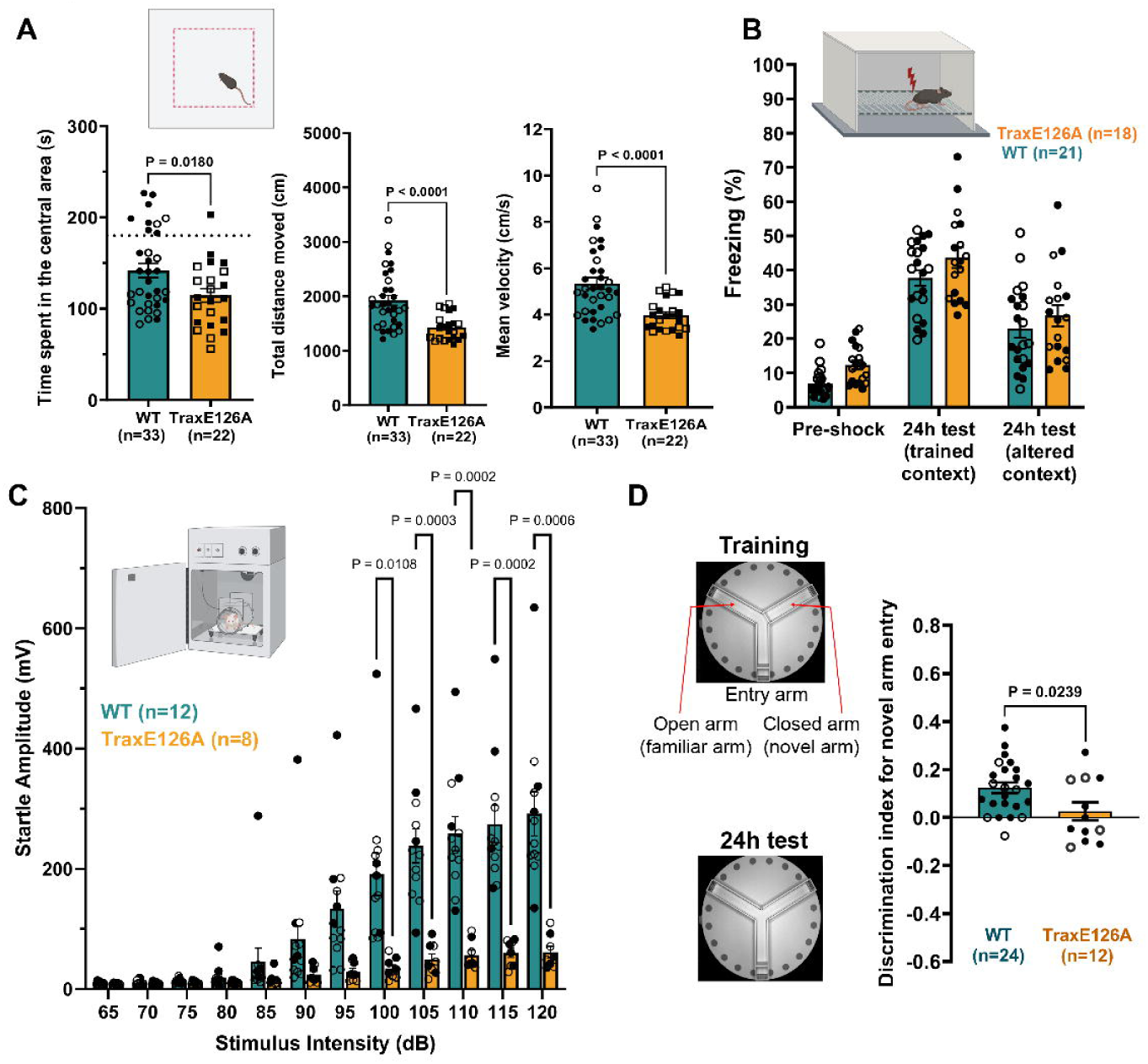
TraxE126A mutant mice show altered open-field behavior, acoustic startle response and impaired long-term memory in spatial Y-maze task but show no deficits in contextual fear memory. **(A)** TraxE126A mutant mice (n=22; 12 males, 10 females) show altered open-field behavior as they spend less time in the central area of the arena (Unpaired t-test, two-tailed, t=2.440, df=53, P=0.018, eta squared = 0.101), travel lesser distance (Welch’s t-test, two-tailed, t=4.733, df=47.96, P<0.0001, eta squared = 0.3184) and travel at lesser mean velocity (Welch’s t-test, two-tailed, t=4.733, df=47.96, P<0.0001, eta squared = 0.3184) compared to WT littermate mice (n=33; 18 males, 15 females). **(B)** TraxE126A mutant mice (n=18; 10 males, 8 females) show no deficits in long-term contextual fear memory compared to WT littermate mice (n=21; 11 males, 10 females) (Two-way repeated measures ANOVA, no main effect of genotype: F (1,37) = 3.730, P=0.061; no interaction between genotype and sessions: F (2, 74) = 0.2297, P=0.795). **(C)** TraxE126A mutant mice show deficits in acoustic startle response compared to WT littermates. The response is significantly dampened in the TraxE126A mutant mice (n=8; 4 males, 4 females) beyond 100dB stimulus intensity compared WT mice (n=12; 4 males, 8 females) (Two-way repeated measures ANOVA, main effect of genotype: F (1,18) = 17.84, P=0.0005; Interaction between stimulus intensity and genotype: F (11, 198) = 19.60, P<0.0001; Sidak’s multiple comparison tests: 100dB, P=0.0108; 105dB, P=0.0003; 110dB, P=0.0002; 115dB, P=0.0002; 120dB, P=0.0006). **(D)** TraxE126A mutant mice show deficits in long-term memory in a spatial 2-trial Y-maze task compared to WT littermates. The WT mice (n=24; 17 males, 7 females) make significantly more entries into the novel arm as shown by their positive discrimination index for the novel arm over familiar arm, whereas TraxE126A mutant mice (n=12; 8 males, 4 females) show significantly lower discrimination for the novel arm (Unpaired t-test, two-tailed, t=2.364, df=34, P=0.0239, eta squared = 0.1412). In all bar graphs, open symbols represent females and closed symbols represent males. The error bars represent SEM in all graphs.

## Discussion

The TN/TX complex has been implicated in multiple cellular processes including processing of miRNAs and pre-miRNAs during RNA interference^23^, hippocampal synaptic plasticity^41,50^, vascular stiffness^45,46^ and adiposity^42^. Additionally, Translin and Trax have been shown to have independent functions in homologous recombination and DNA damage repair^25,75^, cell division and proliferation^29,30^, DNA/RNA metabolism^33–35^, mesenchymal cell differentiation^75,76^, sperm maturation^32^. In this study, we investigated the specific role of the RNase activity of the TN/TX complex in the adult hippocampus using the TraxE126A point mutant mice in which the RNase activity of this complex has been abolished. At the molecular level, we found alterations in the levels of miRNAs in the TraxE126A mice. Forty-three of the predicted targets of these upregulated miRNAs were found to be significantly downregulated at the mRNA level. These genes are involved in focal adhesion, integrin signaling, and extracellular matrix-mediated signaling in the hippocampus. At the synaptic level, we found that TraxE126A mice, which lack the RNase activity of the TN/TX complex, exhibit deficits only in specific forms of hippocampal synaptic plasticity such as spaced 4-train LTP and STC. At the behavioral level, we observed deficits in long-term spatial memory and behavioral alterations in the TraxE126A mice. These findings show for the first time the functional role of TN/TX RNase activity in the brain.

Our data reveal that the TN/TX RNase activity is involved in the regulation of multiple small RNA species including miRNAs, tRNAs and tsRNAs. Inactivating the RNase activity of this complex led to alterations in the levels of many of the small RNAs in the hippocampus of naïve mice. Although our hypothesis was that the TN/TX RNase complex acts to degrade precursor and mature miRNAs, we found miRNA levels being both upregulated and downregulated. This could be due to multiple reasons. First, the mouse model used in this study is a constitutive mutant and it is possible that the alterations in some miRNAs could be a compensatory mechanism during development. Second, changes in some miRNAs could be induced secondary to changes in miRNAs directly targeted by the RNase. Third, TN/TX RNase potentially plays a role in the biogenesis of miRNAs. A direct validation of TN/TX RNase targeting sequences on the miRNAs found to be altered in our study and use of conditional deletion of the RNase activity would be helpful in this regard and this can be explored in future experiments. Nevertheless, among the common miRNAs we identified using both small RNA sequencing and microarray, we found a majority (10 out of 12) to be upregulated. Interestingly, some of these upregulated miRNAs (miR-412, miR-496a, miR-330) were also found to be upregulated in the mouse striatum in a previous study following shRNA-mediated downregulation of Trax^43^. MiR-409, another miRNA found to be upregulated in the TraxE126A mutants, was validated as a target of the TN/TX RNase in an *in vitro* cleavage assay^42^. Additionally, several miRNAs (miR-412, miR-93, miR-7b, miR-1946b) identified to be upregulated in the hippocampus of TraxE126A mice in this study either by sequencing or by microarray were also found to be upregulated in the nucleus accumbens of Translin KO mice in our recent study^44^ while some other miRNAs (mir-409, miR-412, miR-330, miR-370) were found to be upregulated in the cerebellum of Translin KO mice^39^. Given that the RNase activity of the complex is conferred by Trax, and that Translin KO mice lack Trax protein, this provides further support that these miRNAs are targeted for degradation by the TN/TX RNase activity.

A surprising finding from our small RNA microarray analysis is the extensive alteration in the levels of tsRNAs in the hippocampus of TraxE126A mutant mice. Strikingly, almost all the significantly altered tsRNAs were derived from the 5’-end of tRNAs. It is important to note that the TN/TX complex (C3PO) has been previously shown to be involved in tRNA maturation acting as an RNase to remove the 5’ pre-tRNA fragments after the processing of pre-tRNAs by RNase P^34^. Therefore, it is possible that the lack of the RNase activity of TN/TX complex in the E126A mutants leads to the accumulation of the 5’ pre-tRNA-derived fragments. An alternative possibility is that in the absence of the TN/TX RNase activity, other endonucleases cleave the accumulating pre-tRNAs resulting in increased levels of tsRNAs. Significant upregulation of some mature tRNAs was found in the hippocampus of TraxE126A mutant mice, which was also reported in mouse embryonic fibroblasts derived from Translin KO mice^34^ (which also lack Trax protein^52^). Given the evidence that increased levels of 5’-tsRNA fragments can inhibit translational machinery^47,77^, it will be interesting to investigate whether this could contribute to the synaptic plasticity and memory deficits observed in TraxE126A mutants. Evidence is emerging implicating tsRNAs in neuronal physiology and in neurodevelopmental, neurodegenerative, neurobehavioral disorders^78^ as well as metabolic disorders^79^ and uncovering the role of TN/TX RNase in tsRNA biology could potentially open up new therapeutic avenues.

Our results show that abolishing the RNase activity of the TN/TX complex leads to impairments in only selected forms of hippocampal synaptic plasticity, such as spaced-tetanic train-induced LTP and STC, whereas a few other forms of LTP and LTD are not affected. This highlights a specific role of the TN/TX RNase in regulating synaptic plasticity. Impairments in spaced-tetanic train-induced LTP and STC are also reported in the CA1 region of Translin KO mice^41,50^, lending support to the role of TN/TX RNase. Some of the mechanisms shown to support heterosynaptic facilitation of synaptic plasticity in STC include remodeling of actin cytoskeleton and synapse^80,81^, activity-dependent transport and/or translation of local plasticity-related products (PRPs)^73,82,83^, activity-dependent secretion of signaling molecules or cleavage of precursor forms of signaling molecules in the extracellular matrix into their active forms^84–86^ and activity-dependent trafficking of signaling molecules, receptors and ion channels into the synaptic membrane^87–90^. Impairments in LTP and STC observed in the TraxE126A mutants could be due to alterations in one or more of these mechanisms. Although our current study does not delineate the exact mechanism, it offers hints at several possibilities including removal of miRNA-mediated translational repression on key target mRNAs, regulation of tRNA maturation and tsRNA-mediated translational control, regulation of ECM components and signaling through integrin and focal adhesions.

Inactivation of the TN/TX RNase activity also led to alterations in open-field behavior. The point mutant mice spent less time in the central area of the open-field and also moved less. This could indicate more anxiety or decreased motivation for exploration or possibly deficits in motor function. In our previous study, we have shown prominent expression of Translin and Trax in striatal dopaminergic neurons^91^. It is possible that the loss of TN/TX RNase activity in these neurons might impact motor or motivational phenotypes. Future studies aimed at investigating striatal and cerebellar function in these mutants would prove useful in this regard. Translin-null mice, which lack both Translin and Trax proteins, were reported to not show significant alterations in locomotor activity in open-field^51^, although the trend was towards decreased activity. In the contextual fear conditioning task, Trax point-mutant mice showed a trend for marginally higher freezing levels compared to WT mice, further suggesting increased anxiety-like phenotype. Interestingly, enhanced contextual freezing was also reported in female Translin KO mice^51^. However, Stein et al.^51^ found reduced anxiety-related behavior in Translin KO mice in elevated zero maze test and light-dark box exploration test. Trax point-mutant mice showed reduced startle responses to acoustic stimuli over 100dB, similar to as seen in male Translin KO mice^51^. Whether this is partly due to hearing impairments is not investigated in this study. In future experiments, it will be interesting to see if the Trax point mutants show deficits in prepulse inhibition, which is known to be disrupted in several neuropsychiatric disorders^92^. In the two-trial Y-maze task, we found that the Trax point mutant mice showed weaker discrimination for the novel arm, indicating deficits in long-term spatial memory. Although this task has been mostly used to test working memory at shorter intervals, our study shows the utility of two-trial Y-maze task as a long-term memory task. In future studies, we will further characterize this task and utilize in the functional target validation studies. Overall, the behavioral alterations in the TraxE126A mutant mice seem to phenocopy Translin KO mice only in specific phenotypes whereas in some others they differ. There also seems to be sex-specific differences. One explanation for this discrepancy would be that the complete loss of both Trax and Translin proteins in the Translin KO mice leads to the loss of their other functions as well whereas in the TraxE126A mice only the RNase activity of the TN/TX complex is specifically abolished.

For the downstream target prediction and functional enrichment analysis, in this study, we focused only on the miRNAs identified to be significantly altered both in miRNA sequencing and in microarray analysis because of the higher confidence level in these. The predicted targets of these miRNAs were functionally annotated to be involved in multiple cellular processes and signaling pathways. It is certainly possible that other miRNAs that were identified to be significantly altered only in the sequencing approach or in the microarray approach may also regulate important downstream targets. We will investigate these miRNAs and their targets in our future studies. To gain insight into the mRNA targets and signaling pathways impacted by the inactivation of the TN/TX RNase, we performed mRNA sequencing from the same samples which revealed a vast majority of mRNAs to be downregulated, indicating the overall impact of abolished TN/TX RNase activity on gene expression. Functional enrichment analysis of the significantly altered genes showed extracellular matrix-mediated signaling, integrin signaling, focal adhesion and immune system-related pathways to be predominantly impacted. When we further analyzed the genes that were predicted targets of the upregulated miRNAs and were also found to be downregulated in mRNA sequencing, we again found these genes to functionally enrich for focal adhesion, ECM-mediated signaling and integrin signaling.

Our results show downregulation of several ECM components including many collagens, multiple integrin subunits and tenascin-C (Tnc) in the hippocampus of TraxE126A mutants. Components of ECM modulate the function of local receptors or ion channels and send diffuse molecular signals using products of activity-dependent proteolytic cleavage. Because of their role in regulation of synaptic plasticity and remodeling involving both neurons and glia, they have been suggested as components of the ‘tetrapartite synapse’^93,94^. ECM and integrins exhibit extensive crosstalk and are important regulators of several intracellular signaling pathways underlying synaptic plasticity and memory^95^. Tnc, one of the target genes downregulated in the hippocampus of TraxE126A mutants, is an ECM glycoprotein expressed during the development of the nervous system and its expression is upregulated following synaptic stimulation^96,97^. Mice lacking Tnc show impairments in selective forms of synaptic plasticity in hippocampal CA3-CA1 synapses^98–100^, normal spatial learning^98^ but impaired fear memory extinction^99^. Integrin alpha 5 (Itga5) is another target gene downregulated in the hippocampus of TraxE126A mutants. Itga5 regulates spine and synapse formation in hippocampal neurons via Src kinase and Rac signaling^101^. Abnormal structure of perineuronal nets resulting from deficiency of ECM components could lead to subregion-specific changes in perisomatic GABAergic inhibition resulting in metaplastic changes to synaptic plasticity^102,103^. Focal adhesions are transmembrane structures that connect ECM and cytoskeleton to regulate various cellular processes. Integrins are an important part of the signaling through focal adhesions. The ECM-integrin-focal adhesion signaling is capable of relaying both ‘outside-in’ and ‘inside-out’ signaling, thereby acting as a conduit between the interior of the cell and the extracellular environment^104^, which could be affected in the TraxE126A mutants. Our results also reveal the alteration in genes involved in innate immunity and inflammation response. Recent studies have uncovered mechanisms linking innate immunity and long-term memory^105,106^. The study by Jovasevic et al. showed that memory formation is accompanied by increased DNA damage and TLR9-mediated inflammatory signaling^105^. Given the known function of Trax in DNA repair^26,49^, and our results implicating TN/TX RNase in immune response, it will be interesting to investigate whether a combination of these mechanisms might underlie the plasticity and memory deficits in TraxE126A mutant mice. Together our analyses suggest that alterations in these signaling pathways could contribute to the impairments in hippocampal synaptic plasticity and memory in the TraxE126A mutant mice, and we will focus our investigation on these pathways in future studies.

In summary, our study offers a comprehensive analysis of the consequences of abolishing the TN/TX RNase activity on various classes of small RNAs and gene expression in the hippocampus, which consequently leads to impairments in synaptic plasticity, behavior, and long-term memory.

## Author contributions

MSS, JMB and TA designed the study. MSS performed the electrophysiology experiments and hippocampal sample collection, behavioral experiments, and data analysis. JK performed RNA isolation and qRT-PCR experiments. MSS and JK performed RNA sequencing data analysis and functional annotation. XF and JMB generated the TraxE126A mouse model. MCL performed some of the behavioral experiments and helped analyze the open-field data. SMT performed the acoustic startle experiments and analyzed data. TCC helped with some behavioral experiments and colony maintenance. MSS wrote the manuscript with inputs from other authors.

## Acknowledgements

We thank members of the Abel lab and Baraban lab for help with the project and for helpful discussions. We thank Jia Ern Ong, Achala Thippeswamy, Deaven Denis, Jacob Belardo and Quinn Truax for their help with colony maintenance and genotyping in the Abel lab. We thank the Iowa Institute of Human Genomics (IIHG) Core facility for help with miRNA and mRNA sequencing. We thank the Neural Circuits and Behavioral Core (NCBC) at the University of Iowa for the use of their facilities for behavioral experiments.

## Funding sources

This work was supported by funding from the National Institutes of Health R01 MH 087463 to T.A. T.A. is also supported by the Roy J. Carver Charitable Trust.

## Declaration of Interests

The authors declare no competing interests.

## Notes

### Competing Interest Statement

The authors have declared no competing interest.

## References

1 Carthew, R. W. & Sontheimer, E. J. Origins and Mechanisms of miRNAs and siRNAs. Cell 136, 642–655 (2009). 10.1016/j.cell.2009.01.035

2 Fabian, M. R. & Sonenberg, N. The mechanics of miRNA-mediated gene silencing: a look under the hood of miRISC. Nat Struct Mol Biol 19, 586–593 (2012). 10.1038/nsmb.2296

3 Dana, H. et al. Molecular Mechanisms and Biological Functions of siRNA. Int J Biomed Sci 13, 48–57 (2017).

4 Bartel, D. P. MicroRNAs: target recognition and regulatory functions. Cell 136, 215–233 (2009). 10.1016/j.cell.2009.01.002

5 Duchaine, T. F. & Fabian, M. R. Mechanistic Insights into MicroRNA-Mediated Gene Silencing. Cold Spring Harb Perspect Biol 11 (2019). 10.1101/cshperspect.a032771

6 Bartel, D. P. Metazoan MicroRNAs. Cell 173, 20–51 (2018). 10.1016/j.cell.2018.03.006

7 Jungers, C. F. & Djuranovic, S. Modulation of miRISC-Mediated Gene Silencing in Eukaryotes. Front Mol Biosci 9, 832916 (2022). 10.3389/fmolb.2022.832916

8 Rajgor, D. & Hanley, J. G. The Ins and Outs of miRNA-Mediated Gene Silencing during Neuronal Synaptic Plasticity. Noncoding RNA 2 (2016). 10.3390/ncrna2010001

9 Lugli, G., Torvik, V. I., Larson, J. & Smalheiser, N. R. Expression of microRNAs and their precursors in synaptic fractions of adult mouse forebrain. J Neurochem 106, 650–661 (2008). 10.1111/j.1471-4159.2008.05413.x

10 Epple, R. et al. The Coding and Small Non-coding Hippocampal Synaptic RNAome. Mol Neurobiol 58, 2940–2953 (2021). 10.1007/s12035-021-02296-y

11 Steward, O. mRNA localization in neurons: a multipurpose mechanism? Neuron 18, 9–12 (1997). 10.1016/s0896-6273(01)80041-6

12 Schratt, G. microRNAs at the synapse. Nat Rev Neurosci 10, 842–849 (2009). 10.1038/nrn2763

13 Perez, J. D., Fusco, C. M. & Schuman, E. M. A Functional Dissection of the mRNA and Locally Synthesized Protein Population in Neuronal Dendrites and Axons. Annu Rev Genet 55, 183–207 (2021). 10.1146/annurev-genet-030321-054851

14 Baraban, J. M., Shah, A. & Fu, X. Multiple Pathways Mediate MicroRNA Degradation: Focus on the Translin/Trax RNase Complex. Adv Pharmacol 82, 1–20 (2018). 10.1016/bs.apha.2017.08.003

15 Fu, X., Shah, A. & Baraban, J. M. Rapid reversal of translational silencing: Emerging role of microRNA degradation pathways in neuronal plasticity. Neurobiol Learn Mem 133, 225–232 (2016). 10.1016/j.nlm.2016.04.006

16 Cajigas, I. J. et al. The local transcriptome in the synaptic neuropil revealed by deep sequencing and high-resolution imaging. Neuron 74, 453–466 (2012). 10.1016/j.neuron.2012.02.036

17 Glock, C. et al. The translatome of neuronal cell bodies, dendrites, and axons. Proc Natl Acad Sci U S A 118 (2021). 10.1073/pnas.2113929118

18 Hu, Z. & Li, Z. miRNAs in synapse development and synaptic plasticity. Curr Opin Neurobiol 45, 24–31 (2017). 10.1016/j.conb.2017.02.014

19 Buffington, S. A., Huang, W. & Costa-Mattioli, M. Translational control in synaptic plasticity and cognitive dysfunction. Annu Rev Neurosci 37, 17–38 (2014). 10.1146/annurev-neuro-071013-014100

20 Fukao, A., Aoyama, T. & Fujiwara, T. The molecular mechanism of translational control via the communication between the microRNA pathway and RNA-binding proteins. RNA Biol 12, 922–926 (2015). 10.1080/15476286.2015.1073436

21 Koester, S. K. & Dougherty, J. D. A Proposed Role for Interactions between Argonautes, miRISC, and RNA Binding Proteins in the Regulation of Local Translation in Neurons and Glia. J Neurosci 42, 3291–3301 (2022). 10.1523/JNEUROSCI.2391-21.2022

22 Aoki, K., Ishida, R. & Kasai, M. Isolation and characterization of a cDNA encoding a Translin-like protein, TRAX. FEBS Lett 401, 109–112 (1997). 10.1016/s0014-5793(96)01444-5

23 Liu, Y. et al. C3PO, an endoribonuclease that promotes RNAi by facilitating RISC activation. Science 325, 750–753 (2009). 10.1126/science.1176325

24 Ye, X. et al. Structure of C3PO and mechanism of human RISC activation. Nat Struct Mol Biol 18, 650–657 (2011). 10.1038/nsmb.2032

25 Wang, J. Y., Chen, S. Y., Sun, C. N., Chien, T. & Chern, Y. A central role of TRAX in the ATM-mediated DNA repair. Oncogene 35, 1657–1670 (2016). 10.1038/onc.2015.228

26 Chien, T. et al. GSK3beta negatively regulates TRAX, a scaffold protein implicated in mental disorders, for NHEJ-mediated DNA repair in neurons. Mol Psychiatry 23, 2375–2390 (2018). 10.1038/s41380-017-0007-z

27 Aoki, K. et al. A novel gene, Translin, encodes a recombination hotspot binding protein associated with chromosomal translocations. Nat Genet 10, 167–174 (1995). 10.1038/ng0695-167

28 Wu, X. Q., Gu, W., Meng, X. & Hecht, N. B. The RNA-binding protein, TB-RBP, is the mouse homologue of translin, a recombination protein associated with chromosomal translocations. Proc Natl Acad Sci U S A 94, 5640–5645 (1997). 10.1073/pnas.94.11.5640

29 Ishida, R. et al. A role for the octameric ring protein, Translin, in mitotic cell division. FEBS Lett 525, 105–110 (2002). 10.1016/s0014-5793(02)03095-8

30 Yang, S. & Hecht, N. B. Translin associated protein X is essential for cellular proliferation. FEBS Lett 576, 221–225 (2004). 10.1016/j.febslet.2004.08.082

31 Gomez-Escobar, N. et al. Translin facilitates RNA polymerase II dissociation and suppresses genome instability during RNase H2– and Dicer-deficiency. PLoS Genet 18, e1010267 (2022). 10.1371/journal.pgen.1010267

32 Morales, C. R. et al. A TB-RBP and Ter ATPase complex accompanies specific mRNAs from nuclei through the nuclear pores and into intercellular bridges in mouse male germ cells. Dev Biol 246, 480–494 (2002). 10.1006/dbio.2002.0679

33 Jaendling, A. & McFarlane, R. J. Biological roles of translin and translin-associated factor-X: RNA metabolism comes to the fore. Biochem J 429, 225–234 (2010). 10.1042/BJ20100273

34 Li, L. et al. The translin-TRAX complex (C3PO) is a ribonuclease in tRNA processing. Nat Struct Mol Biol 19, 824–830 (2012). 10.1038/nsmb.2337

35 Gupta, A., Pillai, V. S. & Chittela, R. K. Translin: A multifunctional protein involved in nucleic acid metabolism. J Biosci 44 (2019).

36 Sun, C. N. et al. Rescue of p53 blockage by the A(2A) adenosine receptor via a novel interacting protein, translin-associated protein X. Mol Pharmacol 70, 454–466 (2006). 10.1124/mol.105.021261

37 Aisiku, O. R., Runnels, L. W. & Scarlata, S. Identification of a novel binding partner of phospholipase cbeta1: translin-associated factor X. PLoS One 5, e15001 (2010). 10.1371/journal.pone.0015001

38 Tian, Y. et al. Multimeric assembly and biochemical characterization of the Trax-translin endonuclease complex. Nat Struct Mol Biol 18, 658–664 (2011). 10.1038/nsmb.2069

39 Asada, K. et al. Rescuing dicer Defects via Inhibition of an Anti-Dicing Nuclease. Cell Reports 9, 1471–1481 (2014). 10.1016/j.celrep.2014.10.021

40 Li, Z., Wu, Y. & Baraban, J. M. The Translin/Trax RNA binding complex: clues to function in the nervous system. Biochim Biophys Acta 1779, 479–485 (2008). 10.1016/j.bbagrm.2008.03.008

41 Park, A. J. et al. Learning induces the translin/trax RNase complex to express activin receptors for persistent memory. Elife 6 (2017). 10.7554/eLife.27872

42 Fu, X. et al. Genetic inactivation of the translin/trax microRNA-degrading enzyme phenocopies the robust adiposity induced by Translin (Tsn) deletion. Mol Metab 40, 101013 (2020). 10.1016/j.molmet.2020.101013

43 Weng, Y. T. et al. TRAX Provides Neuroprotection for Huntington’s Disease Via Modulating a Novel Subset of MicroRNAs. Mov Disord 37, 2008–2020 (2022). 10.1002/mds.29174

44 Fu, X. P. et al. Translin deletion impairs cocaine-induced locomotor sensitization and RGS8 expression in the nucleus accumbens. Acta Pharmacol Sin (2025). 10.1038/s41401-025-01565-z

45 Tuday, E. et al. Deletion of the microRNA-degrading nuclease, translin/trax, prevents pathogenic vascular stiffness. Am J Physiol Heart Circ Physiol 317, H1116–H1124 (2019). 10.1152/ajpheart.00153.2019

46 Tuday, E. et al. Degradation of Premature-miR-181b by the Translin/Trax RNase Increases Vascular Smooth Muscle Cell Stiffness. Hypertension 78, 831–839 (2021). 10.1161/HYPERTENSIONAHA.120.16690

47 Ivanov, P., Emara, M. M., Villen, J., Gygi, S. P. & Anderson, P. Angiogenin-induced tRNA fragments inhibit translation initiation. Mol Cell 43, 613–623 (2011). 10.1016/j.molcel.2011.06.022

48 Fagan, S. G., Helm, M. & Prehn, J. H. M. tRNA-derived fragments: A new class of non-coding RNA with key roles in nervous system function and dysfunction. Prog Neurobiol 205, 102118 (2021). 10.1016/j.pneurobio.2021.102118

49 Chern, Y. et al. Trax: A versatile signaling protein plays key roles in synaptic plasticity and DNA repair. Neurobiol Learn Mem 159, 46–51 (2019). 10.1016/j.nlm.2018.07.003

50 Park, A. J., Shetty, M. S., Baraban, J. M. & Abel, T. Selective role of the translin/trax RNase complex in hippocampal synaptic plasticity. Mol Brain 13, 145 (2020). 10.1186/s13041-020-00691-5

51 Stein, J. M. et al. Behavioral and neurochemical alterations in mice lacking the RNA-binding protein translin. J Neurosci 26, 2184–2196 (2006). 10.1523/JNEUROSCI.4437-05.2006

52 Chennathukuzhi, V. et al. Mice deficient for testis-brain RNA-binding protein exhibit a coordinate loss of TRAX, reduced fertility, altered gene expression in the brain, and behavioral changes. Mol Cell Biol 23, 6419–6434 (2003). 10.1128/MCB.23.18.6419-6434.2003

53 Thomson, P. A. et al. Association between the TRAX/DISC locus and both bipolar disorder and schizophrenia in the Scottish population. Mol Psychiatry 10, 657–668, 616 (2005). 10.1038/sj.mp.4001669

54 Schosser, A. et al. Association of DISC1 and TSNAX genes and affective disorders in the depression case-control (DeCC) and bipolar affective case-control (BACCS) studies. Mol Psychiatry 15, 844–849 (2010). 10.1038/mp.2009.21

55 Okuda, A. et al. Translin-associated factor X gene (TSNAX) may be associated with female major depressive disorder in the Japanese population. Neuromolecular Med 12, 78–85 (2010). 10.1007/s12017-009-8090-1

56 Weng, Y. T., Chien, T., Kuan, II & Chern, Y. The TRAX, DISC1, and GSK3 complex in mental disorders and therapeutic interventions. J Biomed Sci 25, 71 (2018). 10.1186/s12929-018-0473-x

57 Chennathukuzhi, V. M., Kurihara, Y., Bray, J. D., Yang, J. & Hecht, N. B. Altering the GTP binding site of the DNA/RNA-binding protein, Translin/TB-RBP, decreases RNA binding and may create a dominant negative phenotype. Nucleic Acids Res 29, 4433–4440 (2001). 10.1093/nar/29.21.4433

58 Martin, M. Cutadapt removes adapter sequences from high-throughput sequencing reads. EMBnet.journal 17, 10–12 (2011). doi: 10.14806/ej.17.1.200.

59 Langmead, B. Aligning short sequencing reads with Bowtie. Curr Protoc Bioinformatics Chapter 11, Unit 11 17 (2010). 10.1002/0471250953.bi1107s32

60 Love, M. I., Huber, W. & Anders, S. Moderated estimation of fold change and dispersion for RNA-seq data with DESeq2. Genome Biol 15, 550 (2014). 10.1186/s13059-014-0550-8

61 Chen, Y. & Wang, X. miRDB: an online database for prediction of functional microRNA targets. Nucleic Acids Res 48, D127–D131 (2020). 10.1093/nar/gkz757

62 Dobin, A. et al. STAR: ultrafast universal RNA-seq aligner. Bioinformatics 29, 15–21 (2013). 10.1093/bioinformatics/bts635

63 Huang da, W., Sherman, B. T. & Lempicki, R. A. Systematic and integrative analysis of large gene lists using DAVID bioinformatics resources. Nat Protoc 4, 44–57 (2009). 10.1038/nprot.2008.211

64 Sherman, B. T. et al. DAVID: a web server for functional enrichment analysis and functional annotation of gene lists (2021 update). Nucleic Acids Res 50, W216–W221 (2022). 10.1093/nar/gkac194

65 Shetty, M. S. et al. Mice Lacking the cAMP Effector Protein POPDC1 Show Enhanced Hippocampal Synaptic Plasticity. Cereb Cortex 32, 3457–3471 (2022). 10.1093/cercor/bhab426

66 Walsh, E. N., Shetty, M. S., Diba, K. & Abel, T. Chemogenetic Enhancement of cAMP Signaling Renders Hippocampal Synaptic Plasticity Resilient to the Impact of Acute Sleep Deprivation. eNeuro 10 (2023). 10.1523/ENEURO.0380-22.2022

67 Gould, T. J. et al. Sensorimotor gating deficits in transgenic mice expressing a constitutively active form of Gs alpha. Neuropsychopharmacology 29, 494–501 (2004). 10.1038/sj.npp.1300309

68 Kelly, M. P. et al. Developmental etiology for neuroanatomical and cognitive deficits in mice overexpressing Galphas, a G-protein subunit genetically linked to schizophrenia. Mol Psychiatry 14, 398–415, 347 (2009). 10.1038/mp.2008.124

69 Friard, O. & Gamba, M. BORIS: a free, versatile open-source event-logging software for video/audio coding and live observations. Methods in Ecology and Evolution 7, 1325–1330 (2016). 10.1111/2041-210X.12584

70 Ewald, J., Zhou, G., Lu, Y. & Xia, J. Using ExpressAnalyst for Comprehensive Gene Expression Analysis in Model and Non-Model Organisms. Curr Protoc 3, e922 (2023). 10.1002/cpz1.922

71 Akiyoshi, K. et al. Adenosine A(2A) Receptor Regulates microRNA-181b Expression in Aorta: Therapeutic Implications for Large-Artery Stiffness. J Am Heart Assoc 12, e028421 (2023). 10.1161/JAHA.122.028421

72 Frey, U. & Morris, R. G. Synaptic tagging and long-term potentiation. Nature 385, 533–536 (1997). 10.1038/385533a0

73 Redondo, R. L. & Morris, R. G. Making memories last: the synaptic tagging and capture hypothesis. Nat Rev Neurosci 12, 17–30 (2011). 10.1038/nrn2963

74 Okuda, K., Hojgaard, K., Privitera, L., Bayraktar, G. & Takeuchi, T. Initial memory consolidation and the synaptic tagging and capture hypothesis. Eur J Neurosci 54, 6826–6849 (2021). 10.1111/ejn.14902

75 Kasai, M. et al. The translin ring specifically recognizes DNA ends at recombination hot spots in the human genome. J Biol Chem 272, 11402–11407 (1997). 10.1074/jbc.272.17.11402

76 Ikeuchi, Y. et al. Translin modulates mesenchymal cell proliferation and differentiation in mice. Biochem Biophys Res Commun 504, 115–122 (2018). 10.1016/j.bbrc.2018.08.141

77 Shi, J., Zhang, Y., Zhou, T. & Chen, Q. tsRNAs: The Swiss Army Knife for Translational Regulation. Trends Biochem Sci 44, 185–189 (2019). 10.1016/j.tibs.2018.09.007

78 Tian, H., Hu, Z. & Wang, C. The Therapeutic Potential of tRNA-derived Small RNAs in Neurodegenerative Disorders. Aging Dis 13, 389–401 (2022). 10.14336/AD.2021.0903

79 Zong, T. et al. tsRNAs: Novel small molecules from cell function and regulatory mechanism to therapeutic targets. Cell Prolif 54, e12977 (2021). 10.1111/cpr.12977

80 Ramachandran, B. & Frey, J. U. Interfering with the actin network and its effect on long-term potentiation and synaptic tagging in hippocampal CA1 neurons in slices in vitro. J Neurosci 29, 12167–12173 (2009). 10.1523/JNEUROSCI.2045-09.2009

81 Pinho, J., Marcut, C. & Fonseca, R. Actin remodeling, the synaptic tag and the maintenance of synaptic plasticity. IUBMB Life 72, 577–589 (2020). 10.1002/iub.2261

82 Doyle, M. & Kiebler, M. A. Mechanisms of dendritic mRNA transport and its role in synaptic tagging. EMBO J 30, 3540–3552 (2011). 10.1038/emboj.2011.278

83 Okada, D., Ozawa, F. & Inokuchi, K. Input-specific spine entry of soma-derived Vesl-1S protein conforms to synaptic tagging. Science 324, 904–909 (2009). 10.1126/science.1171498

84 Ishikawa, Y., Horii, Y., Tamura, H. & Shiosaka, S. Neuropsin (KLK8)-dependent and –independent synaptic tagging in the Schaffer-collateral pathway of mouse hippocampus. J Neurosci 28, 843–849 (2008). 10.1523/JNEUROSCI.4397-07.2008

85 Lu, Y. et al. TrkB as a potential synaptic and behavioral tag. J Neurosci 31, 11762–11771 (2011). 10.1523/JNEUROSCI.2707-11.2011

86 Pang, P. T. & Lu, B. Regulation of late-phase LTP and long-term memory in normal and aging hippocampus: role of secreted proteins tPA and BDNF. Ageing Res Rev 3, 407–430 (2004). 10.1016/j.arr.2004.07.002

87 Okuno, H. et al. Inverse synaptic tagging of inactive synapses via dynamic interaction of Arc/Arg3.1 with CaMKIIbeta. Cell 149, 886–898 (2012). 10.1016/j.cell.2012.02.062

88 Park, A. J. et al. A presynaptic role for PKA in synaptic tagging and memory. Neurobiol Learn Mem 114, 101–112 (2014). 10.1016/j.nlm.2014.05.005

89 Park, P. et al. On the Role of Calcium-Permeable AMPARs in Long-Term Potentiation and Synaptic Tagging in the Rodent Hippocampus. Front Synaptic Neurosci 11, 4 (2019). 10.3389/fnsyn.2019.00004

90 Yao, Y. et al. PKM zeta maintains late long-term potentiation by N-ethylmaleimide-sensitive factor/GluR2-dependent trafficking of postsynaptic AMPA receptors. J Neurosci 28, 7820–7827 (2008). 10.1523/JNEUROSCI.0223-08.2008

91 Fu, X. et al. Elevated body fat increases amphetamine accumulation in brain: evidence from genetic and diet-induced forms of adiposity. Transl Psychiatry 11, 427 (2021). 10.1038/s41398-021-01547-9

92 Swerdlow, N. R., Braff, D. L. & Geyer, M. A. Sensorimotor gating of the startle reflex: what we said 25 years ago, what has happened since then, and what comes next. J Psychopharmacol 30, 1072–1081 (2016). 10.1177/0269881116661075

93 Dityatev, A. & Rusakov, D. A. Molecular signals of plasticity at the tetrapartite synapse. Curr Opin Neurobiol 21, 353–359 (2011). 10.1016/j.conb.2010.12.006

94 Dityatev, A., Seidenbecher, C. I. & Schachner, M. Compartmentalization from the outside: the extracellular matrix and functional microdomains in the brain. Trends Neurosci 33, 503–512 (2010). 10.1016/j.tins.2010.08.003

95 Park, Y. K. & Goda, Y. Integrins in synapse regulation. Nat Rev Neurosci 17, 745–756 (2016). 10.1038/nrn.2016.138

96 Dityatev, A. & Schachner, M. Extracellular matrix molecules and synaptic plasticity. Nat Rev Neurosci 4, 456–468 (2003). 10.1038/nrn1115

97 Nakic, M., Manahan-Vaughan, D., Reymann, K. G. & Schachner, M. Long-term potentiation in vivo increases rat hippocampal tenascin-C expression. J Neurobiol 37, 393–404 (1998).

98 Evers, M. R. et al. Impairment of L-type Ca2+ channel-dependent forms of hippocampal synaptic plasticity in mice deficient in the extracellular matrix glycoprotein tenascin-C. J Neurosci 22, 7177–7194 (2002). 10.1523/JNEUROSCI.22-16-07177.2002

99 Morellini, F. et al. Impaired Fear Extinction Due to a Deficit in Ca(2+) Influx Through L-Type Voltage-Gated Ca(2+) Channels in Mice Deficient for Tenascin-C. Front Integr Neurosci 11, 16 (2017). 10.3389/fnint.2017.00016

100 Strekalova, T. et al. Fibronectin domains of extracellular matrix molecule tenascin-C modulate hippocampal learning and synaptic plasticity. Mol Cell Neurosci 21, 173–187 (2002). 10.1006/mcne.2002.1172

101 Webb, D. J., Zhang, H., Majumdar, D. & Horwitz, A. F. alpha5 integrin signaling regulates the formation of spines and synapses in hippocampal neurons. J Biol Chem 282, 6929–6935 (2007). 10.1074/jbc.M610981200

102 Saghatelyan, A. K. et al. Reduced Perisomatic Inhibition, Increased Excitatory Transmission, and Impaired Long-Term Potentiation in Mice Deficient for the Extracellular Matrix Glycoprotein Tenascin-R. Molecular and Cellular Neuroscience 17, 226–240 (2001). 10.1006/mcne.2000.0922

103 Bukalo, O., Schachner, M. & Dityatev, A. Hippocampal Metaplasticity Induced by Deficiency in the Extracellular Matrix Glycoprotein Tenascin-R. The Journal of Neuroscience 27, 6019–6028 (2007). 10.1523/jneurosci.1022-07.2007

104 Chastney, M. R., Conway, J. R. W. & Ivaska, J. Integrin adhesion complexes. Curr Biol 31, R536–R542 (2021). 10.1016/j.cub.2021.01.038

105 Jovasevic, V. et al. Formation of memory assemblies through the DNA-sensing TLR9 pathway. Nature 628, 145–153 (2024). 10.1038/s41586-024-07220-7

106 Shen, Y. et al. CCR5 closes the temporal window for memory linking. Nature 606, 146–152 (2022). 10.1038/s41586-022-04783-1

