## Supplementary Figures and Tables for "Genetic inactivation of the Translin/Trax RNase activity alters small RNAs including miRNAs, disrupts gene expression and impairs distinct forms of hippocampal synaptic plasticity and memory"

**Supplementary Figure S1.** qRT-PCR analysis of some selected mRNAs predicted as targets of the differentially expressed miRNAs on miRDB database. Data are plotted as fold change from WT (normalized to housekeeping control HPRT levels).

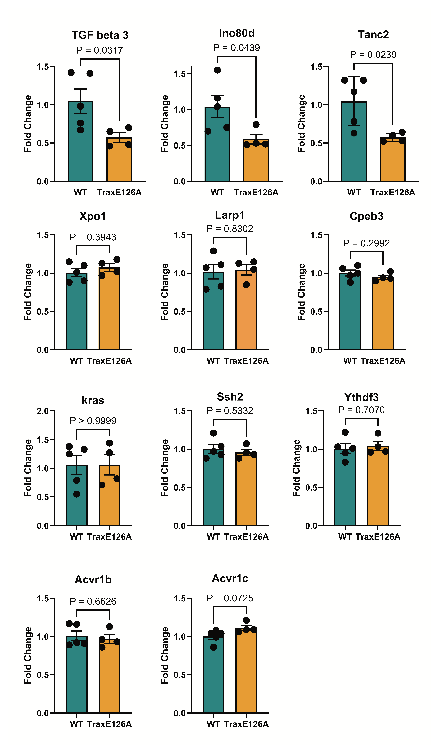

**Supplementary Table T1.** Primer pairs used for qPCR validation

| **Target** | **Forward primer** | **Reverse primer** |
| --- | --- | --- |
| Tgfb3 | 5’-GGCCCTGGACACCAATTACT-3’ | 5’-GGTTCGTGGACCCATTTCCA-3’ |
| Tanc2 | 5’-CTCTGGGCAAGGTGGCAT-3’ | 5’-CAGATGGAGGTGGGCCAAG-3’ |
| Ino80d | 5’-TAGCCCCTCCTACAGGGTTC-3’ | 5’-AGGGGTGTCCCTCTAATCCC-3’ |
| Xpo1 | 5’-AGTGCCTCACTGAGATTGCTGG-3’ | 5’-CCCATTCGAGTATGCAAGTCGG-3’ |
| Larp1 | 5’-ACTCCATGCTTTGGAGGGTG-3’ | 5’-AGGTATGGGAGCCTCTTGGA-3’ |
| Cpeb3 | 5’-TGGCACTTAACACCAGGAGC-3’ | 5’-TGGCTGTCATCCAAGAAGGC-3’ |
| Kras | 5’-AGCCAGGAGTCAAGGTGAAA-3’ | 5’-GTCAAGGCGCTCTTGCCTAC-3’ |
| Ssh2 | 5’-TTGGATGGTGATGGTGGGTT-3’ | 5’-CAAGCCTTGTGCAAACTCTGT-3’ |
| Ythdf3 | 5’-GGTGGATAGCTGTTATTCTGATTTG-3’ | 5’-GGTGGATAGCTGTTATTCTGATTTG-3’ |
| Acvr1b | 5’-GAACCGCTACACAGTGACCA-3’ | 5’-AATTCCCGGCTTCCCTTGAG-3’ |
| Acvr1c | 5’-CACAGTTCCAACAACGTGACC-3’ | 5’-CAAGAGAGGCAGACCTGTGG-3’ |
| Tnc | 5’-GAGACCTGACACGGAGTATGAG-3’ | 5’-CTCCAAGGTGATGCTGTTGTCTG-3’ |
| Itga5 | 5’-TCCGTGGAGTTTTACCGGC-3’ | 5’-CAACCAGCACACTGACTCTTTC-3’ |
| Plaur | 5’-AGGACTACCGTGCTTCGGGAAT-3’ | 5’-ACACGGTCTCTGTCAGGCTGAT-3’ |
| Pxn | 5’-GTGAGAAGGACTACCACAGCCT-3’ | 5’-GGACCAAAGAAGGCTCCACACT-3’ |
| Tln1 | 5’-AGTGACGGACAGCATCAACCAG-3’ | 5’-GGATTCTCCAGGAGTTCTCGGA-3’ |
| Hprt | 5’-TTGCTGACCTGCTGGATTACA-3’ | 5’-CCCCGTTGACTGATGATTACA-3’ |

**Supplementary Figure S2.** TraxE126A mutant mice show no alterations in basal synaptic transmission and very short-term plasticity. **(A)** Basal field-EPSP amplitudes are not significantly different between wildtype mice and TraxE126A mutant mice (two-way repeated measures ANOVA; no main effect of genotype, F(1,40) = 2.134; P = 0.1519; no interaction). **(B)** Presynaptic fiber volley (PFV) amplitudes are not significantly different between wildtype mice and TraxE126A mutant mice (two-way repeated measures ANOVA; no main effect of genotype, F(1,40) = 0.1964; P = 0.660; no interaction). **(C)** Paired-pulse facilitation of field-EPSP slopes over a range of interstimulus intervals is not significantly different between wildtype mice and TraxE126A mutant mice (two-way repeated measures ANOVA; no main effect of genotype, F(1,30) = 2.177; P = 0.1505; no interaction).

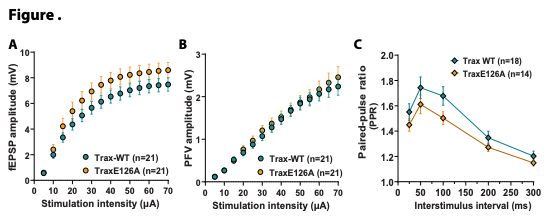

**Supplementary Figure S3.** In the 2-trial Y-maze task for long-term memory assessment, the discrimination index for the novel arm exploration time is not significantly different between TraxE126A mutants (n=12; 8 males, 4 females) and WT mice (n=24; 17 males, 7 females) (Unpaired t-test, two-tailed, t=0.442, df=34, P=0.661, eta squared = 0.0057). Discrimination Index (DI) = (Time exploring novel arm – Time exploring familiar arm) / Total time exploring both arms). In the bar graph, open symbols represent females and closed symbols represent males.

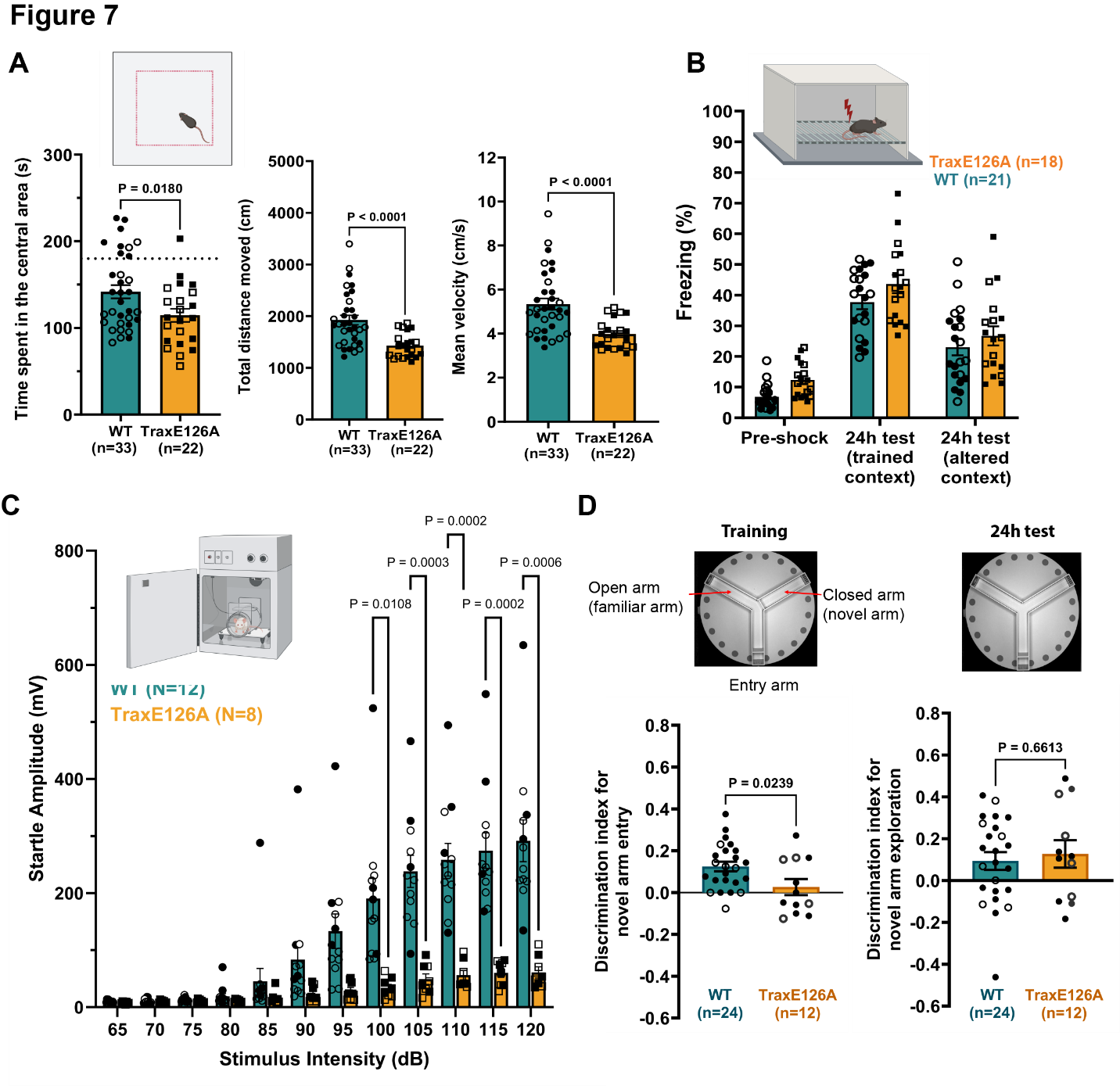

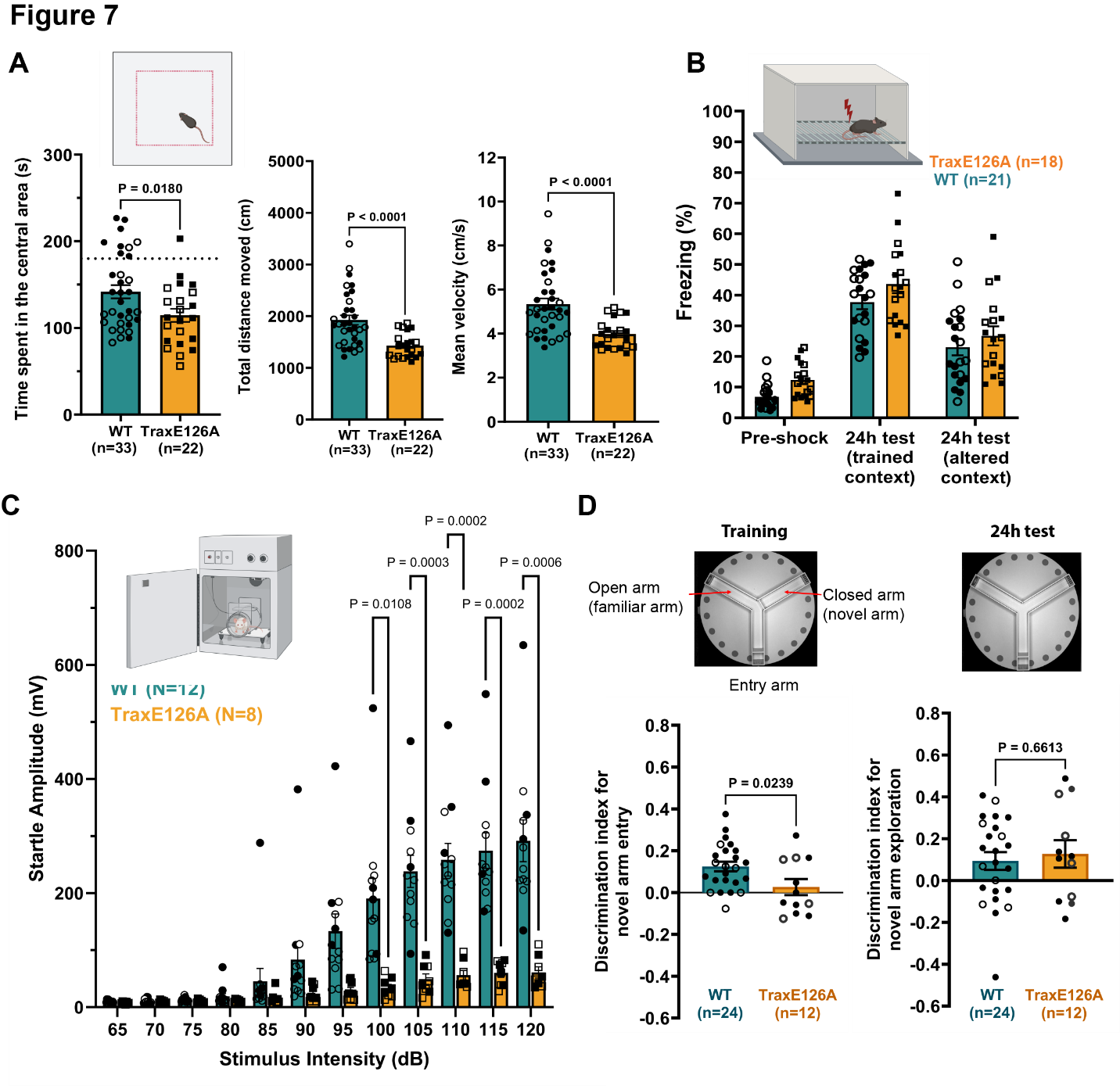

**Supplementary Figure S4.** Scoring criteria in the 2-trial Y-maze task. The videos of test session were manually scored using the open-source BORIS software. For number of entries into a given arm, an event was considered an 'arm entry' if both the front paws of the mouse are past the 'blocker slit' for the arm being scored (blue lines in the schematic below). For the exploration time of the novel or familiar arm, time spent exploring the left or right arm including the area on the respective side of the vertical red line in the schematic below was scored (this was because the mice can sit outside the arm and sniff the entry sites without fully entering). Any time spent grooming was not scored as exploration. Time spent in the entry arm was not scored.

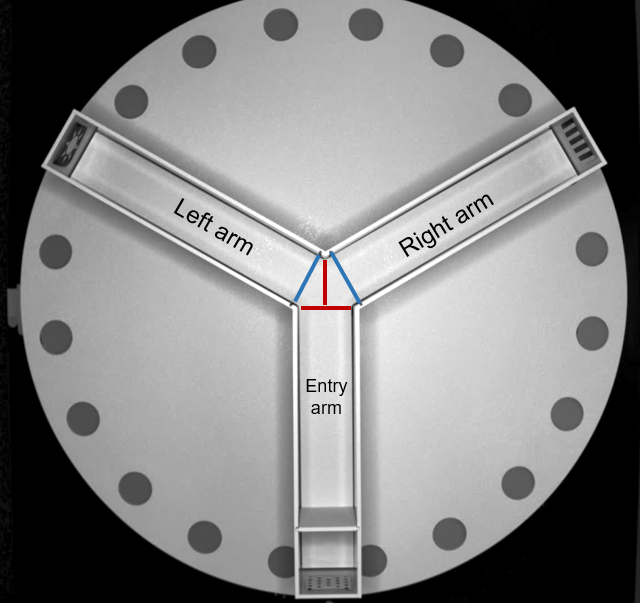
